# Neurodegeneration and neuroinflammation are linked, but independent of **α**-synuclein inclusions, in a seeding/spreading mouse model of Parkinson’s disease

**DOI:** 10.1101/2020.08.05.237750

**Authors:** Pierre Garcia, Wiebke Jürgens-Wemheuer, Oihane Uriarte, Kristopher J Schmit, Annette Masuch, Simone Brioschi, Andreas Weihofen, Eric Koncina, Djalil Coowar, Tony Heurtaux, Enrico Glaab, Rudi Balling, Carole Sousa, Alessandro Michelucci, Tony Kaoma, Nathalie Nicot, Tatjana Pfander, Walter Schulz-Schaeffer, Ahmad Allouche, Nicolas Fischer, Knut Biber, Michel Mittelbronn, Manuel Buttini

## Abstract

A key process of neurodegeneration in Parkinson’s disease (PD) is the transneuronal spreading of α-synuclein. Alpha-synuclein is a presynaptic protein that is implicated in the pathogenesis of PD and other synucleinopathies, where it forms, upon intracellular aggregation, pathological inclusions. Other hallmarks of PD include neurodegeneration and microgliosis in susceptible brain regions. Whether it is primarily transneuronal spreading of α-synuclein particles, inclusion formation, or other mechanisms, such as inflammation, that cause neurodegeneration in PD is unclear. We used spreading/aggregation of α-synuclein induced by intracerebral injection of α-synuclein preformed fibrils into the mouse brain to address this question. We performed quantitative histological analysis for α-synuclein inclusions, neurodegeneration, and microgliosis in different brain regions, and a gene expression profiling of the ventral midbrain, at two different timepoints after disease induction. We observed significant neurodegeneration and microgliosis in brain regions not only with, but also without α-synuclein inclusions. We also observed prominent microgliosis in injured brain regions that did not correlate with neurodegeneration nor with inclusion load. In longitudinal gene expression profiling experiments, we observed early and unique alterations linked to microglial mediated inflammation that preceded neurodegeneration, indicating an active role of microglia in inducing neurodegeneration. Our observations indicate that α-synuclein inclusion formation is not the major driver in the early phases of PD-like neurodegeneration, but that diffusible, oligomeric α-synuclein species, which induce unusual microglial reactivity, play a key role in this process. Our findings uncover new features of α-synuclein induced pathologies, in particular microgliosis, and point to the necessity of a broader view of the process of “prion-like spreading” of that protein.

## Introduction

Protein misfolding and aggregation are central pathological processes in neurodegenerative diseases, where they are believed to play a key role in driving the pathology [1, 2]. Proteins such as the amyloid beta peptide (Aβ) and tau in Alzheimer’s disease (AD), TAR DNA-binding protein 43 (TDP43) in fronto-temporal dementia or motor neuron disease, prion in Creutzfeldt-Jakob disease, and finally alpha-synuclein (α-syn) in Parkinson’s disease (PD), are all examples of physiologically occurring proteins that, upon pathological misfolding, form oligomers, fibrils, and extracellular (Aβ, prion) or intracellular (TDP43, tau, α-syn) deposits, and injure neurons in the process [3–5].

An important property of these disease-associated proteins is their ability to self-propagate. This has been known for decades in prion diseases, in which disease-associated misfolding proteins themselves are sufficient to induce the disease process when transposed into a susceptible recipient host already decades ago [3, 6]. They do so by acting as a seed and corrupting the endogenous form of the protein, leading it to aggregate and form, over time, intracellular inclusions along interconnected neuronal pathways [7, 8]. Currently, the predominant narrative of this “spreading hypothesis” is that misfolded/aggregated particles of a disease protein move transsynaptically from neuron to neuron, causing dysfunction and damage along the way [4, 8]. Major support for this hypothesis comes from two observations. First, the neuropathological studies by Braak and colleagues, staging tau inclusions in AD [9], and Lewy inclusions in PD [10], suggest a progressive appearance, starting first in a population of susceptible neurons, of proteinaceous intraneuronal inclusions, a process that takes place over decades. Second, postmortem studies of in PD patients that had received intrastriatal fetal neuron transplants to combat dopamine loss, revealed Lewy bodies in a subset of the grafted neurons, indicating a spreading of abnormal α-syn from the diseased neurons of the recipient to those of the donor [11, 12].

Alpha-syn is a presynaptic protein that normally is involved in the regulation of the synaptic vesicle cycle [13, 14]. Its involvement in PD was discovered when it was identified as an essential component of a PD pathological hallmark, the Lewy body [15], and when mutations in its gene, as well as dupli-or triplication thereof, were shown to lead to hereditary forms of the disease [16, 17]. Its prion-like spreading properties have been demonstrated in *in vitro* and *in vivo* model systems, using intracranial injection of Lewy-body containing brain extracts, of viral-construct mediated α-syn overexpression and/or administration of pre-formed fibrils (PFFs) from recombinant α-syn as seeds to induce spreading and aggregation[18] [19–22].

The role of α-syn spreading and inclusion formation is PD pathogenesis still unclear, since no correlation between PD symptoms and α-syn inclusion load was consistently found [23–26]. Different possibilities that could explain what ultimately causes neuronal dysfunction and injury and, hence, neurological symptoms, have recently surfaced [27, 28]. Among those, the notion that smaller moieties, or oligomers, of misfolded proteins rather than more fibrillar, deposited forms of α-syn are the most neurotoxic has gained traction [29, 30]. It is important to note that, in the field of AD, it was thought for decades that the Aβ deposited into plaques is the most harmful to neurons, whereas more recent evidence points to diffusible, soluble small Aβ moieties as being the major neurotoxic form [31]. In the PD field though, this notion is still debated: while some studies report the potential harm α-syn oligomers can cause [30, 32], other studies still contend that neuronal dysfunction and injury cannot occur without the demonstrable presence of inclusions [33, 34].

In this study, we addressed this issue in the using a α-syn seeding/spreading induced in wildtype mice [20]. We used intracranial administration of recombinant murine α-syn PFFs to induce α-syn spreading and inclusion formation in the brain of wildtype mice, and examined neurodegeneration and microgliosis in brain regions with α-syn inclusions and, importantly, in those without. We observed neurodegeneration in both cases, indicating that neuronal injury can occur independently of the progressive formation of α-syn inclusions. Because neuroinflammation has emerged as a key player in neurodegenerative disease [35], and because microglia are the main cellular effectors of this process [36, 37], we also measured microgliosis in our model. We noticed, in regions with or without inclusions, a surprisingly strong microgliosis (4-5x over baseline), which far surpassed that observed after administration of neurotoxins such as the dopaminergic lesioning agent 6-hydroxydopamine (6-OHDA). In contrast to mice injected with 6-OHDA, neurodegeneration and microgliosis did not correlate with each other in the brains of α-syn PFFs injected mice. Moreover, by measuring gene expression profiles after intrastriatal α-syn PFF injection, we observed numerous significant changes in inflammation-related genes and pathways, and an unusual microglial molecular activation profile that preceded neuron loss and indicated a direct involvement of these cells in the neurodegeneration process.

These findings indicated that, in this model, microgliosis does not occur primarily as a response to neuronal damage, but is likely part of an intrinsic response to a process that is independent of the progression of α-syn inclusion formation. Our results demonstrate that PD-like neurodegeneration can occur independently of the presence of α-syn inclusions, and thus that PD-like pathologies is more than just the progressive formation of pathological inclusions. It may involve spreading of other, more soluble forms of toxic aggregates, such as oligomers, inducing an excessive microglial response, before inclusions are formed. We believe these results add an important aspect on how the pathogenic properties of “prion-like” α-syn should be viewed, and how future therapeutic interventions for PD will be designed.

## Materials and Methods

### Expression, and purification of recombinant murine α-Syn, and generation of pre-formed fibrils (PFFs), and of oligomers

Expression and purification of recombinant murine α-syn and generation of PFFs were performed as described [38]. PFFs were stored aliquoted at −80°C until use. For the preparation of oligomers, recombinant α-syn was purchased from Analytik Jena (Jena, Germany). Oligomers were generated as described [39, 40] by incubating soluble α-syn in 10 mM Tris-HCl, 100 mM NaCl under continuous shaking in an Eppendorf Thermomixer at 650rpm and 37°C for 24 h, then stored aliquoted at 2 mg/ml at −80°C until use.

### Western Blot of α-syn PFFs and oligomers

The composition of α-syn PFFs and oligomers was checked by non-denaturing Western Blot. Three different concentrations of oligomers of PFFs (10 ng, 100 ng, 500 ng), were loaded on 4-10% Precast Gel Mini Protean TGX (BioRad) according to manufacturer’s instructions. To reveal α-syn bands, anti-synuclein antibody clone 4D6 (Covance) was used at 1:2000 dilution (2 h at RT incubation), followed by IRDye^R^ 800 CW donkey anti-mouse, diluted 1:10.000 (1 h at RT incubation). Image was captured with a LI-COR Bioscience C-Digit Chemoluminescence scanner.

### Animals

Three- to 6-month-old C57Bl/6J mice were purchased from Jackson via Charles River (Bois-des-Oncins, France), or Janvier Labs (Le-Genet-St.-Isle, France). Mice were housed in individually-ventilated cages (IVC) in a conventional animal facility of the University of Luxembourg, or in the facility of SynAging, in Vandeouvre-les-Nancy, France. All animal studies were in agreement with the requirements of the EU Directive 2010/63/EU and Commission recommendation 2007/526/EC. Male or female mice were housed under a 12h-12h dark/light cycle with *ad libitum* access to water and food (#2016, Harlan, Horst, NL). For time point of 90 dpi PFF injections (see below), the youngest mice were used, for time point 13 dpi PFF injections, the oldest mice were used, so that, at euthanasia, all the mice were of comparable age (6 to 6.5 months). Otherwise, mice were assigned randomly to study groups. For quantitative histology (see below), 10-11 mice/group were used, and all were quantified, whereas for transcriptional profiling, 6 mice/group were used. Such numbers have proven sufficient in previous studies on different models of neurodegeneration, while also keeping in line with the rule of the “3Rs” [41–45]. Animal studies were approved by the institutional Animal Experimentation Ethics Committee of the University of Luxembourg and the responsible Luxembourg government authorities (Ministry of Health, Ministry of Agriculture). Alternatively, experiments done at the SynAging site were approved by ethics committee “Comité d’Ethique Lorrain en Matière d’Expérimentation Animale”, and by the governmental agency the “Direction Départementale de la Protection des Populations de Meurthe et Moselle-Domaine Expérimentation Animale”.

### Intrastriatal injections of α-syn PFFs, α-syn oligomers, and 6-hydroxydopamine

Alpha-syn PFFs were sonicated in a sonicating waterbath (Branson 2510, Danbury, CT) for 2 hours at RT, keeping the temperature constant at 25°C by adding ice as needed, or using the Bioruptor UCD 300 (Diagenode, Seraing, Belgium) with 30 cycles of 15sec ON/ 15sec OFF at 4°C. Sonicated PFFs were kept on ice and used within ten hours. Mice were injected under isoflurane anesthesia (2%) on a heating pad. A 1 cm long mid-line scalp incision was made into the desinfected surgical area and a 0.5 mm hole drilled unilaterally into the skull using stereotaxic coordinates for striatum according to the Mouse Brain Atlas of Franklin and Paxinos [46]. Ten µg of PFFs, or respective PBS solution (control mice) were administered, in volumes of 2 μl, within the right dorsal striatum at the following relative-to-bregma coordinates: anterior +0.5 mm, lateral +2.1 mm; depth +3.2 mm. The 24-gauge blunt tip needle of the Hamilton syringue (7105KH, Bonaduz, CH) was inserted down 3.3 mm for 10 seconds to form an injection pocket, and the needle remained in place for 2 minutes before and after the injection procedure. The hole was covered with bonewax (Lukens, Arlington, VA) and the wound closed using 7mm Reflex wound clips (Fine Science Tools, Heidelberg, Germany). Two % xylocaine gel was applied to the wound, and mice were allowed to recover from anesthesia before being put back into their home cages. The day of injection of PFFs was named day 0. Same coordinates and similar procedure were used for 6-OHDA or α-syn oligomers. Striatal injection of 6-OHDA has been described elsewehere [47]. Striatal injections of α-syn oligomers were done with 4 μg oligomers in 2 μl vehicle. Control mice received the same volume of vehicle (see above). Mice were euthanized in a deep anaesthesia (i.p. injection of Medetomidin, 1mg/kg and Ketamin, 100mg/kg) by transcardial transfusion with PBS. PFF-injected mice were euthanized either at day 13 (13 dpi) or at day 90 (90 dpi) after intrastriatal injections (“day 0”: day of injection). Mice injected with oligomers or with 6-OHDA were euthanised at 13 dpi.

### Tissue extraction and preparation

For immunohistochemistry, extracted brains were fixed in in 4% buffered PFA for 48h and kept in PBS with 0.1% NaN3 until they were cut with a vibratome (VT1000 S from Leica) into sagittal 50µm free-floating sections. Before the staining procedure, sections were kept at −20°C in a cryoprotectant medium (1:1 v/v PBS/Ethylene Glycol, 10g.L^-1^ Polyvinyl Pyrrolidone). Alternatively, for dopamine measurement or RNA extraction, after removal from the skull, brains were dissected on ice into regions, and isolated striatum and ventral midbrain were quickly weighted, then snap-frozen on dry ice until further processing. Extraction and measurement of striatal dopamine (DA) has been described elsewhere [48]. Briefly, after homogenization and derivatization, striatal metabolites were measured with a gas-chromatography/mass-spectrometry set-up (Agilent 7890B GC – Agilent 5977A MSD, Santa Clara, CA). Absolute level of DA were determined using an internal standard, 2-(3,4-Dihydroxyphenyl)ethyl-1,1,2,2-d_4_-amine HCl (D-1540, C/D/N isotopes, Pointe-Claire, Canada). For RNA extraction from the ventral midbrain, the RNEasy Universal Kit (Quiagen) was used, after homogenization of midbrain tissues in a Retsch MM 400 device (2 min at 22Hz, Haan, Germany). RNA concentrations and integrity were determined using a Nanodrop 2000c (Thermo Scientific) and a BioAnalyzer 2100 (Agilent), respectively. Purified RNAs were considered of sufficient quality if their RNA Integrity Number (RIN) was above 8.5, their 260/230 absorbance ratio > 1, and their 260/280 absorbance ratio = 2.

### Single and double-label immunohistochemistry

Immunostaining procures followed standard protocols, as described [41, 42]. All stainings, except those for proteinase-K resistant α-syn inclusions (see below), were performed on free-floating 50 μm-thick sections. Supplemental Table 1 lists all primary and secondary antibodies used in this study, as well as their dilutions. All other reagents were from Sigma unless indicated otherwise. All antibody incubations were at room temperature, except for the anti-synaptophysin antibody, which was incubated at 4°C. Sections were washed 3 x in PBS between each incubation step. To block endogenous peroxidases and for permeabilization, sections were incubated with 3% H_2_O_2_ v/v and 1.5% Triton X100 v/v for 30 min. For immoperoxidase staining with anti-synuclein antibody, this step was followed by an epitope unmasking step with 75% v/v formic acid for 5 min. To avoid unspecific antibody binding, sections were incubated with 5% serum (Vector Laboratories, Burlingame, CA) or 5% BSA w/v in PBS for 1h before they were incubated with the respective primary antibody, or antibodies in case of double labeling. The following day, sections were incubated with a secondary antibody for 1-2 hours (fluorophore-coupled for immunofluorescence, or biotinylated for immunoperoxidase). Singly or doubly fluorescently-stained sections were mounted on Superfrost plus slides (Thermoscientific, Walham, MA), air-dried, and coverslipped using ProLong Gold antifade mountant (Life technologies, Darmstadt, Germany). For immunoperoxidase staining, antibody binding was visualized using an ABC Vectastain Kit (Vector Laboratories), followed by detection with diaminobenzidine (Merck) and H_2_O_2_ as peroxidase substrates. Sections were mounted, dried overnight and coverslipped with Neo-mount (Merck) after soaking in Neo-clear xylene substitute (Merck) for 10 min. Visualization of Proteinase-K resistant α-synuclein inclusions was done by paraffin-embedded tissue blot (PET blot) on 3µm paraffin sections mounted on nitrocellulose membrane (0.45µm, BioRad) as previously [49].

### Proximity ligation assay

Protocol for PLA was adapted to be performed on free floating sections. All reactants were prepared according manufacturer’s recommendations (Duolink. Sigma) and incubation times were as described [50]. Washes were performed in 24-well plates at RT, and reactions volumes were 40µL at 37°C. First, 20µg of anti-pSER129-α-syn mouse monoclonal 11E5 antibody (Prothena Biosciences, see suppl. table 1) were conjugated with either plus or minus oligonucleotide probes according manufacturer’s recommendations, and stored 4°C until use. Free floating sections were washed in PBS and permeabilized as described above. Blocking was performed with DuoLink blocking solution for 2h RT. Sections were incubated overnight with both plus and minus probe-linked antibody (1:1 1/750 in Duolink antibody diluent solution). For ligation of the probes, after washing of the probe-linked antibodies (2 × 5min in Duolink’s Buffer A), the ligation-ligase solution was added and incubated for 30 min at 37°C. For detection, after washing of the ligation-ligase solution (2 × 5min in Duolink’s Buffer A), sections were incubated with the amplification-polymerase solution for 2.5h at 37°C. Sections were then washed in Buffer B for ten minutes, and in Buffer B 0.01X for one minute prior mounting, dried in the dark, and coverslipped. Z-stacks of pictures were acquired at 40X with a Zeiss LabA1 microscope, a maximum intensity projection was created using the Zen Blue 2012 software (Zeiss).

**Table 1:** Unique molecular features of microglial response to intrastriatal α-syn PFF injection. Table shows genes and their coded proteins for factors typically associated with microglial pro-inflammatory M1 profile (highlighted in yellow), for factors typically associated with microglial anti-inflammatory M2 profile (highlighted in green), and for generic microglial activation factors (highlighted in blue). Comparisons are between ventral midbrains of ipsi PFF *versus* ipsi PBS and of ipsi PFF *versus* contra PFF. The rank p-values and the FDRs are given. The proteins whose genes have either a significant p-value (<0.05) or pfp < 0.1 are highlighted in bold. Already at 13 dpi, *Cybb*, the gene for NADPH oxidase 2, a mediator of oxidative stress, was upregulated in ipsi PFF midbrain, a region that showed neurodegeneration at 90 dpi (see above). Another gene, *Ptgs2*, whose product, cyclooxygenase 2, has been reported to generate neurotoxic arachidonic acid metabolites, was also upregulated in ipsi PFF midbrains, most strongly so (pfp <0.1) when comparing ipsi PFF with contra PFF. This gene can be expressed also in astrocytes. *Cd68*, whose protein is linked to lysomal function and possibly acts as a scavenger receptor, was also strongly upregulated in ipsi PFF midbrains. Finally, a triad of factors associated with microglia activation (*Tyrobp, Trem2, Tlr2*) was upregulated at 13 dpi in ipsi PFF midbrains. Their downregulation at 90 dpi in ipsi PFF versus ipsi PBS could indicate that the inflammation process starts to resolve.

| Gene Symbol | Protein | ipsi PFF versus ipsi PBS |  |  |  |  |  | ipsi PFF versus contra PFF |  |  |  |  |  |
| --- | --- | --- | --- | --- | --- | --- | --- | --- | --- | --- | --- | --- | --- |
|  |  | 13dpi |  |  | 90dpi |  |  | 13dpi |  |  | 90dpi |  |  |
|  |  | FC | p-Value | Pfp | FC | p-Value | Pfp | FC | p-Value | FDR | FC | p-Value | Pfp |
| *Cybb* | NAPDH oxidase 2 | 1.407 | 5.17E-05 | 2.24E-02 | -1.221 | 5.36E-03 | 3.95E-01 | 1.413 | 1.09E-04 | 2.06E-02 | 1.155 | 7.08E-02 | 7.60E-01 |
| *Ptgs2* | Cyclooxygenase 2 | 1.285 | 1.50E-03 | 1.74E-01 | -1.189 | 1.18E-03 | 2.06E-01 | 1.506 | 1.28E-05 | 4.52E-03 | 1.436 | 8.06E-05 | 1.84E-02 |
| *Cxcl10* | Chemokine (C-X-C motif) ligand 10 | 1.226 | 6.11E-03 | 3.62E-01 | -1.141 | 1.03E-01 | 9.28E-01 | 1.344 | 7.64E-04 | 7.62E-02 | -1.003 | 5.15E-01 | 1.10E+00 |
| *Cd86* | CD86 antigen | 1.112 | 8.32E-02 | 9.65E-01 | -1.139 | 5.96E-02 | 8.08E-01 | 1.304 | 2.08E-03 | 1.38E-01 | 1.073 | 1.87E-01 | 9.36E-01 |
| *Il1b* | Interleukin 1 beta | 1.131 | 8.25E-02 | 9.62E-01 | 1.071 | 3.38E-01 | 1.04E+00 | 1.085 | 2.54E-01 | 1.06E+00 | 1.091 | 2.41E-01 | 9.72E-01 |
| *Il6* | Interleukin 6 | 1.030 | 4.26E-01 | 1.07E+00 | -1.022 | 4.75E-01 | 1.07E+00 | 1.032 | 4.48E-01 | 1.09E+00 | -1.050 | 3.33E-01 | 1.06E+00 |
| *Tnf* | Tumor necrosis factor alpha | -1.003 | 6.35E-01 | 1.02E+00 | 1.002 | 6.81E-01 | 9.97E-01 | 1.017 | 4.93E-01 | 1.08E+00 | 1.082 | 3.04E-01 | 1.00E+00 |
| *Ccl2* | Chemokine (C-C motif) ligand 2 | -1.023 | 3.86E-01 | 1.07E+00 | -1.073 | 1.92E-01 | 1.03E+00 | -1.050 | 2.35E-01 | 1.06E+00 | 1.082 | 1.46E-01 | 9.10E-01 |
| *Nos2* | Nitric oxide synthase 2, inducible | 1.069 | 2.86E-01 | 1.10E+00 | -1.063 | 3.21E-01 | 1.06E+00 | -1.014 | 6.54E-01 | 1.05E+00 | -1.003 | 6.12E-01 | 1.10E+00 |
| *Il12b* | Interleukin 12b | 1.003 | 6.24E-01 | 1.03E+00 | -1.023 | 5.26E-01 | 1.06E+00 | -1.006 | 6.11E-01 | 1.06E+00 | 1.074 | 3.31E-01 | 1.01E+00 |
| *Il12a* | Interleukin 12a | -1.042 | 4.72E-01 | 1.06E+00 | 1.060 | 2.96E-01 | 1.05E+00 | -1.007 | 6.61E-01 | 1.05E+00 | 1.164 | 8.29E-02 | 7.89E-01 |
| *Mrc1* | Mannose receptor, C type 1 | -1.029 | 2.09E-01 | 1.09E+00 | -1.064 | 3.72E-02 | 7.06E-01 | 1.070 | 1.50E-01 | 9.79E-01 | 1.439 | 5.63E-05 | 1.42E-02 |
| *Tgfb1* | Transforming growth factor, beta 1 | 1.098 | 9.24E-02 | 9.90E-01 | -1.085 | 9.92E-02 | 9.16E-01 | 1.196 | 2.36E-02 | 5.14E-01 | 1.093 | 1.39E-01 | 9.00E-01 |
| *Socs3* | Suppressor of cytokine signaling 3 | 1.037 | 4.87E-01 | 1.06E+00 | -1.014 | 5.17E-01 | 1.06E+00 | 1.031 | 5.01E-01 | 1.08E+00 | -1.029 | 2.80E-01 | 1.04E+00 |
| *Arg1* | Arginase 1 | -1.017 | 4.81E-01 | 1.06E+00 | -1.012 | 5.62E-01 | 1.06E+00 | -1.059 | 3.59E-01 | 1.09E+00 | -1.091 | 1.86E-01 | 9.64E-01 |
| *Il10* | Interleukin 10 | 1.026 | 4.36E-01 | 1.07E+00 | -1.109 | 1.19E-01 | 9.53E-01 | 1.029 | 5.00E-01 | 1.08E+00 | -1.064 | 2.52E-01 | 1.02E+00 |
| *Chil3* | Chitinase-like 3 | 1.058 | 2.19E-01 | 1.09E+00 | -1.100 | 1.29E-01 | 9.62E-01 | 1.066 | 2.42E-01 | 1.06E+00 | -1.023 | 3.25E-01 | 1.06E+00 |
| *Il4* | Interleukin 4 | -1.110 | 1.71E-01 | 1.07E+00 | 1.088 | 2.62E-01 | 1.06E+00 | -1.004 | 6.62E-01 | 1.05E+00 | 1.099 | 2.39E-01 | 9.72E-01 |
| *Retnla* | Resistin like alpha | -1.003 | 3.26E-01 | 1.08E+00 | 1.010 | 4.66E-01 | 1.02E+00 | -1.101 | 1.38E-01 | 9.77E-01 | 1.140 | 1.00E-01 | 8.33E-01 |
| *Cd68* | CD68 antigen | 1.683 | 6.74E-08 | 2.07E-04 | -1.201 | 1.08E-02 | 4.96E-01 | 2.042 | 3.49E-11 | 1.31E-07 | 1.292 | 1.49E-03 | 1.28E-01 |
| *Tyrobp* | TYRO protein tyrosine kinase binding protein | 1.628 | 1.75E-07 | 3.48E-04 | -1.237 | 1.22E-02 | 5.08E-01 | 1.570 | 2.45E-06 | 1.26E-03 | 1.129 | 8.70E-02 | 7.97E-01 |
| *Trem2* | Triggering receptor expressed on myeloid cells 2 | 1.363 | 8.16E-05 | 2.93E-02 | -1.183 | 2.46E-02 | 6.34E-01 | 1.582 | 2.06E-06 | 1.16E-03 | 1.187 | 2.56E-02 | 5.52E-01 |
| *Tlr2* | Toll-like receptor 2 | 1.199 | 1.51E-02 | 5.48E-01 | -1.056 | 2.77E-01 | 1.06E+00 | 1.196 | 2.15E-02 | 4.89E-01 | 1.057 | 2.35E-01 | 9.69E-01 |
| *P2ry6* | Pyrimidinergic receptor P2Y, 6 | 1.273 | 2.68E-03 | 2.45E-01 | -1.144 | 8.74E-02 | 8.97E-01 | 1.477 | 1.64E-05 | 5.53E-03 | 1.064 | 3.39E-01 | 1.02E+00 |
| *Aif1* | Allograft inflammatory factor 1 (Iba1) | 1.167 | 2.21E-02 | 6.37E-01 | -1.097 | 1.53E-01 | 9.92E-01 | 1.225 | 9.88E-03 | 3.31E-01 | 1.147 | 7.10E-02 | 7.61E-01 |
| *Ltb4r1* | Leukotriene B4 receptor 1 | -1.015 | 6.22E-01 | 1.02E+00 | -1.007 | 6.29E-01 | 1.05E+00 | 1.011 | 5.67E-01 | 1.08E+00 | -1.031 | 5.07E-01 | 1.10E+00 |
| *Tmem119* | Transmembrane protein 119 | 1.060 | 3.79E-01 | 1.08E+00 | -1.176 | 5.15E-02 | 7.79E-01 | 1.119 | 1.50E-01 | 9.79E-01 | 1.070 | 3.27E-01 | 1.01E+00 |
| *Siglec1* | Sialic acid binding Ig-like lectin 1 | 1.096 | 2.04E-01 | 1.09E+00 | -1.102 | 1.31E-01 | 9.66E-01 | 1.145 | 9.39E-02 | 8.74E-01 | 1.061 | 4.23E-01 | 1.04E+00 |

### Quantitative neuropathology on immunostained sections

Imaging of peroxidase-labelled sections for pSER129-α-syn, and of fluorescently labelled sections for Tyrosine-Hydroxylase (TH), Dopamine Transporter (DAT), or Ionized calcium binding adaptor molecule 1 (Iba1), was done on a Zeiss LabA1 microscope, coupled to a Zeiss Axiocam MRm3 digital camera, and to a PC using the Zeiss Zen Blue 2012 software.

Alpha-syn inclusions were visualized by immunostaining for pSER129-α-syn. For the quantitation of α-syn inclusions in the frontal cortex and the amygdala (basolateral nucleus), 2 immunoperoxidase labeled (see above) sections/animal were imaged, using the 10x objective (frontal cortex) or the 20x objective (amygdala). A total of 4-6 images were collected for each region (10x objective, 2 x 1.52 mm each image), and digitized. After manually drawing regions of interests and thresholding, the percent image area occupied by immunopositive structures was determined using the ImageJ v. 1.45 (NIH, Bethesda, MD) public domain software. All values obtained from sections of the same animal were averaged. For the quantitation of α-syn inclusions in the SN, double immunostainings for TH and pSER129-α-syn were performed. TH staining was used to locate the SN, and images were acquired at 10x magnification. Percent overlap of pSER129-α-syn signal within the TH immunopositive area was calculated using ImageJ. For the quantitation of synaptophysin-positive synaptic terminals, fluorescently stained sections from each animal were viewed by a Zeiss LSM 710 laser-scanning confocal microscope, using a 20× objective and a software magnification zoom factor was used to obtain images of 180 × 180 μm each. For each animal, a total of 4-6 images were collected from the frontal cortex, and 4 from the hippocampal pyramidal region. Images were then transferred to a PC personal computer, and average intensity of positive presynaptic terminals was quantified for each image using the ImageJ. Values from individual animals were averaged. This method to quantify synaptic integrity has been validated by electron microscope quantitation of synaptic densities in a previous study [41].

The quantitation of degeneration of TH positive neurons in the SN has been described, and results obtained with this approach have been shown to correlate with stereological cell counts (supplemental material in Ashrafi et al. (2017) [47]). For the quantitation of striatal TH-positive neuronal fibers and of DAT-positive synaptic terminal, doubly labelled sections with anti-TH and anti-DAT were used. A total of six to nine 40x pictures (223.8 x 167.7 μm each) of the dorsal striatum, from 2-3 sections per mouse, were acquired using the optical sectioning system Apotome.2 (Zeiss). The percent area occupied by TH and DAT was determined using Image J software and averaged for each mouse.

For the quantitation of microglial activation in the hippocampus and frontal cortex, 2 sections /animal were labeled for the microglial marker *Iba1.* For the frontal cortex, a total of 6, and, for the hippocampus, a total of 3-4 images were collected with a 40x objective (223.8 x 167.7 μm each image). Digitized images were transferred to a laptop computer, and, with ImageJ v. 1.45, after thresholding, average area occupied by Iba1-positive microglia was measured. All values obtained from sections of the same animal were averaged. For the quantitation of the microglial activation in the SN, TH and Iba1-double-labeled sections were imaged with a 10x objective. Average area covered by TH-positive neurons in control mice was used to determine the region of interest, restricted to the SN, to measure microglial activation. Four subregions of the SN were imaged and quantified for each mouse [47]. Iba1 immunopositive cells were quantified within each subregions, averaged for each of them, and converted in mm^2^. For each mouse, the sum of the four averaged subregions was used as a measure for microglial activation.

All quantitative neuropathological analyses were performed blinded on coded sections, and, for each of the measurements, codes were only broken when quantification for that measure in all animals was complete. For all measures, the ipsilateral and contralateral values of PBS-injected control mice were similar (no statistical difference detected), this these values were grouped. Statistics on quantitative histological data were done using the GraphPad Prism 8 software. ANOVA followed by Dunnett’s post-hoc was used for normally distributed data sets. Pearson’s test, or Spearman’s rank test where appropriate, were used to study linear correlations. P values smaller than 5% were considered as significant.

### Microarray analysis and calculation of differentially expressed genes

GeneChip Mouse Gene 2.0ST Arrays (Affymetrix) were used for transcriptional profiling. Total RNAs (150ng) were processed using the Affymetrix GeneChip® WT PLUS Reagent Kit according to the manufacturer’s instructions (Manual Target Preparation for GeneChip® Whole Transcript (WT) Expression Arrays P/N 703174 Rev. 2). In this procedure, adapted from Bougnaud et al. (2016) [51], the purified, sense-strand cDNA is fragmented by uracil-DNA glycosylase (UDG) and apurinic/apyrimidinic endonuclease 1 (APE 1) at the unnatural dUTP residues and breaks the DNA strand. The fragmented cDNA was labelled by terminal deoxynucleotidyl transferase (TdT) using the Affymetrix proprietary DNA Labelling Reagent that is covalently linked to biotin; 5.5 μg of single-stranded cDNA are required for fragmentation and labelling, then 3.5 μg of labeled DNA + hybridization controls were injected into an Affymetrix cartridge. Microarrays were then incubated in the Affymetrix Oven with rotation at 60 rpm for 16 hr at 45°C, then the arrays were washed and scanned with the Affymetrix® GeneChip® Scanner 3000, based on the following protocol: UserGuide GeneChip® Expression Wash, Stain and Scan for Cartridge Arrays P/N 702731 Rev. 4, which generated the Affymetrix raw data CEL files containing hybridization raw signal intensities were imported into the Partek GS software. First, probe intensities were summarized to gene expression signals using Partek default options (GCcontent adjustment, RMA background correction, quantile normalization, log2 transformation and summarization by means).

For statistical analysis, the normalized and log2 transformed data was loaded into the R/Bioconductor statistical environemnt. The rank product (Package: ***RankProd***) approach was chosen to determine the differentially expressed genes (DEGs) [52–54]. Rank product statistics were computed, since they have been shown to enable a robust non-parametric analysis of microarray datasets with limited number of samples [52]. Estimated p-values and pfp (percentage of false prediction) values were determined and used as nomimal and adjusted significance scores, respectively. Pfp scores estimate the significance of differential expression after adjusting for multiple hypothesis testing, and can have values larger than 1. The chosen significance cut-offs were p-value < 0.05 and pfp < 0.1. A relaxed cut-off (pfp < 0.1 instead of < 0.05) was chosen to avoid loss of information for the subsequent enrichment analysis, which combines several genes below this threshold to enable detection of pathway alterations. No minimal fold change threshold was applied.

For visualization of differential gene expression, Venn diagrams and heatmaps were generated using the ***VennDiagram*** and ***gplots*** packages, respectively, in R. Data pre-processing included removal of all transcripts missing gene IDs and duplicated entries (after ranking). To avoid mismatches or other discrepencies, the Affymetrix probe sets IDs were used to calculate the overlapping DEGs between the different comparisons. Diagrams were generated for the following criteria and comparisons: 1) 13 dpi & 90 dpi – ipsiPFF *versus* ipsiPBS (p-value < 0.05); 2) 13 dpi & 90 dpi – ipsiPFF *versus* ipsiPBS (pfp < 0.1); 3) 13 dpi & 90 dpi – ipsiPFF *versus* contraPFF (p-value < 0.05); 4) 13 dpi & 90 dpi – ipsiPFF vs contraPFF (pfp < 0.1). In a second step, we were interested in investigating the expression direction of the overlapping transcripts between the early to late timepoint. Therefore, we extracted the probe IDs, matched the individual lists and grouped them into high and low expressed transcripts. Then, the above mentioned venn diagrams were generated with these newly generated lists. Heatmaps were generated using the ***heatmap.2*** function for ipsiPFF vs ipsiPBS and ipsiPFF vs contraPFF for p-value < 0.05 and pfp < 0.1 at 13 dpi and 90 dpi respectively. The log2-transformed data matrix was used to plot. Additionally, we applied hierarchical top down clustering (***cor*** and ***hclust*** basic R functions) and the data matrix was scaled row-by-row generating Z-scores.

### Gene Set Enrichment Analysis

The enrichment analysis for GO terms (biological processes (BP) only) was performed using the GUI (graphical user interface) version GSEA (version 3.0) published by the Broad Institute (dowload: http://software.broadinstitute.org/gsea/downloads.jsp) [55]. All parameters were set to default in GSEA, except “Collapse dataset to gene symbols” was set to “false”, “Permutation type” was set to “gene_set” and “Max size: excluding larger sets” was set to “250”. One optimization step was introduced: a costumized GMT/GMX file was generated in R/Bioconductor with mouse NCBI EntryzIDs as gene identifiers. This file was used as the “Gene sets database” in GSEA. The resulting enrichment scores (ES) were obtained applying the weighted Kolmogorov-Smirnov-like statistics. ES reflect the level to which a gene set is overrepresented among the top up- or down-regulated genes in a ranked gene list, then, the ES statistic was normalized (normalized enrichment scores, NES) as described [55]. Finally, the p-value significance scores were adjusted for multiple hypothesis testing using the method by Benjamini and Hochberg[56] to provide final FDR scores. A network map of the enrichment analysis results was generated using Cytoscape [57]. The mapping parameters used in Cytoscape were: p-value < 0.05, FDR Q-value < 0.1 (default setting is 1) and Overlap > 0.5. The enrichment map was automatically launched from GSEA and created in Cytoscape. In the enrichment maps, nodes represent enriched gene sets associated with BPs, and edges the degree of similarity between them using the overlap coefficient (threshold > 0.5). Further curation of gene sets was done manually. Since gene sets with similar gene compositions tend to group together, such gene set clusters were easily identifiable. Nodes grouped into more than one gene cluster according to this procedure were assigned to the most overlapping cluster, i.e. the cluster they were associated with by a shorter sequence of connecting edges in the ontology graph. All softwares used are listed in Supplemental table 2.

### Identification of cellular source of DEGs

For the identification of the cellular source of specific DEGs, the public database https://www.brainrnaseq.org/ was used.

## Results

### Western blot characterisation of α-syn moieties

A non-denaturing blot of the α-syn moieties used in this study is shown in supplemental Fig, 1. For intrastriatal injections, α-syn oligomers were used non-sonicated, whereas α-syn PFFs were sonicated. Based on their profile on WB, the oligomer preparation was composed mainly of monomers, dimers, and trimers, as well as higher molecular weight species. The sonicated PFFs were composed mainly of monomers and dimers, and higher molecular weight species. When compared to their non-sonicated counterparts, sonicated PFFs seemed to have less of all these components, consistent with a shearing effect of sonication that produces smaller α-syn fragments, which may then act as seeds.

### Intrastriatal injection of PFFs causes bilateral α-syn inclusions in multiple brain regions

Because we wanted to capture the early features of α-syn spreading associated pathologies, we decided to focus our investigations on time points when, these pathologies have not reached a peak yet [20]. Since α-syn inclusions have been suggested to be a major driver of PD-like pathology [34, 58], we first looked at the appearance of these inclusions in our model.

To determine if intrastriatal injection of murine α-syn PFF reliably induced propagation of fibrillar α-syn in our mice, we first performed immunohistochemistry against pSER129- α-syn on sections of both brain hemispheres 13 days and 90 days after they had been injected with PFFs (13 dpi, 90 dpi). Immunostaining for pSER129- α-syn is the most commonly used approach to detect α-syn inclusions in rodent or human brain tissues [59].

At an early time point after α-syn PFF administration (13 dpi), we only detected few pSER129- α-syn positive inclusions in frontal cortex, amygdala, and SN, and more in the ipsilateral striatum (supplemental Fig. 2).

However, at 90dpi, we observed robust appearance of pSER129- α-syn positive cellular and neuritic inclusions in the ipsi- and contralaterally, in the same brain regions (Fig.1 A). Quantitation of image area occupied revealed median coverage of 10% for the ipsilateral frontal cortex, 5% for the contralateral frontal cortex, 8% for the ipsilateral amygdala, 4.2% for the contralateral amygdala, and 12% for the ipsilateral SN. No or very few α-syn inclusions were found in the contralateral SN and striatum, and no inclusions in either side of the hippocampus. Cells containing inclusions had neuronal morphology. In the ipsilateral SN, fluorescent double staining for pSER129-α-syn and TH, a marker for dopaminergic neurons in the SN, showed that 85% of inclusions co-localized with TH-positive neurons, indicating that most, if not all, inclusions are localized in neurons.

**Figure 1.**
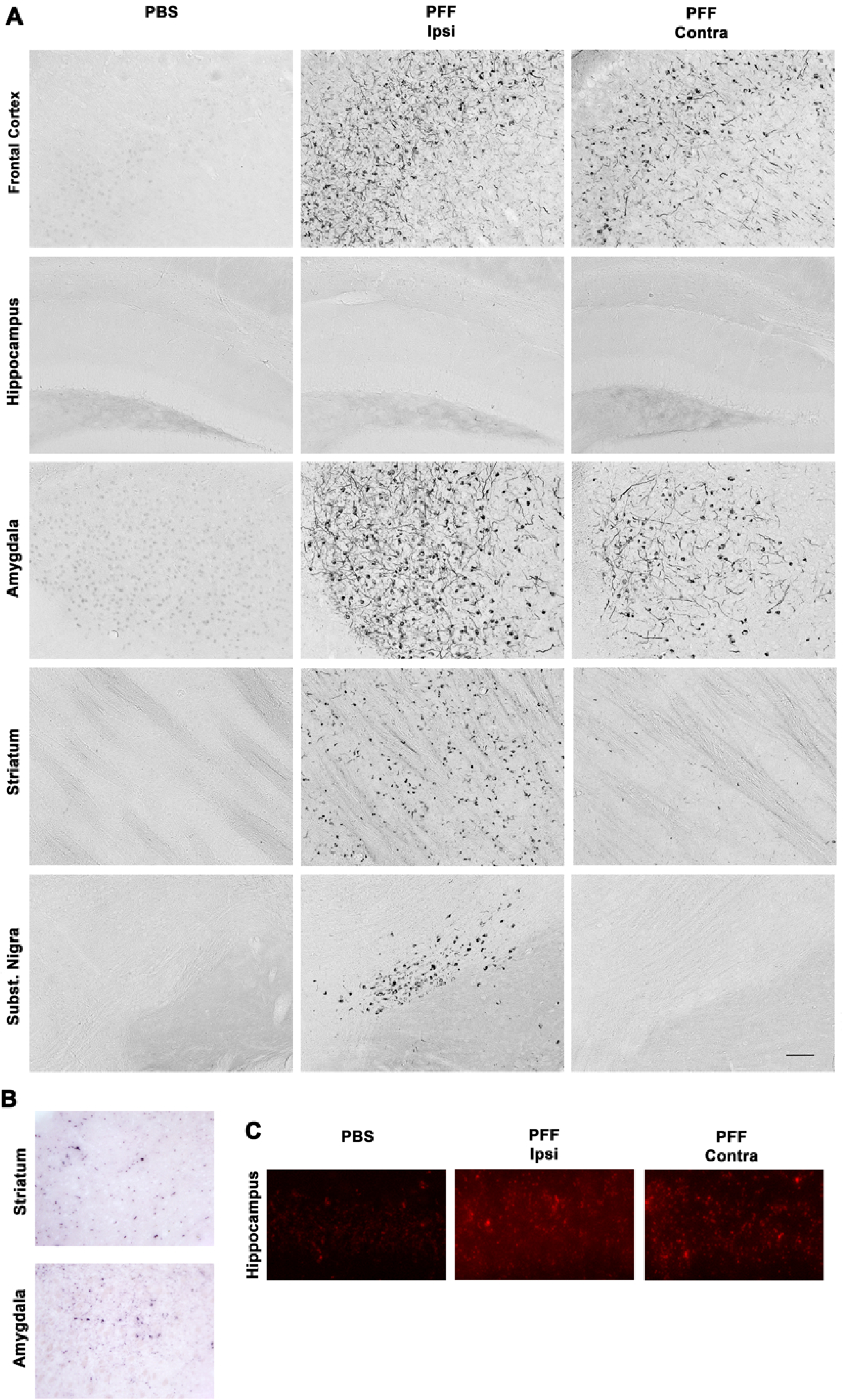
Intrastriatal injection of murine α-syn PFFs induced α-syn inclusions in various brain regions. Mice were euthanized 90 days after injection (90 dpi, n=10-11/group). A. PhosphoSER129 α-syn immunostaining showed numerous α-syn inclusions in neuritic and neuronal body structures in different brain regions. Widespread α-syn inclusions were observed bilaterally in frontal cortex and the amygdala, ipsilaterally in the striatum and the Substantia Nigra (SN), and only minimally in the contralateral striatum and SN. None were observed in the hippocampus. No inclusions were observed in either side of the brains of PBS-injected control mice. Pictures show the ipsilateral side of these mice. B. Proteinase-K digestion on thin sections generated form paraffin-embedded tissue blocks revealed the presence of digestion-resistant α-syn inclusions stained for PhosphoSER129 α-syn. Shown here are ipsilateral striatum and amygdala for illustration. C. Proximity-ligation assay using a monoclonal PhosphoSER129 α-syn antibody showed the presence of enhanced levels of oligomeric forms of α-syn in the hippocampus of PFF-injected mice, where no inclusions could be detected 90 dpi, compared to PBS-injected controls. Scale bar = 250 μm (A), 250 μm (B), 25 μm (C).

To determine if the pSER129- α-syn positive inclusions were Proteniase-K resistant, we performed a Proteinase-K assay PET assay [49]. We observed numerous pSER129- α-syn positive signals in these tissue sections (Fig. 1 B), indicating that most inclusions were proteinase-K resistant.

Inclusions are not the only α-syn species that have been suggested to be linked to neurodegeneration in PD. To determine if regions without detectable α-syn inclusions, such as the hippocampus, were still affected by abnormal α-syn after injection of PFFs, we performed a Proximity Ligation Assay (PLA). This assay has been used for detecting oligomeric forms of α-syn in human [60] and mouse models of PD [61]. We observed greatly enhanced signal intensity in the hippocampi of PFF-injected mice than in those of control mice (Fig. 1C), indicating the presence of abnormal levels of oligomeric α-syn in that area.

Overall, the pattern of α-syn inclusions we observed 90 dpi matches that described at a similar time point by Luk et al. (2012) [20]. The robust appearance of intracellular α-syn inclusions in this model opened up the possibility of analyzing how they are associated with other pathological hallmarks, such as neurodegeneration and –inflammation, and which of these events might precede the others.

### Intrastriatal injection of PFFs causes bilateral synaptic loss and unilateral dopaminergic neuron injury that is independent of **α**-syn inclusions

We set out to determine to what extent the presence of neuronal α-syn deposition was linked to neurodegeneration, 90 dpi after intrastriatal administration of PFFs.

First, we analyzed synaptic degeneration in the hippocampus and frontal cortex. In these brain regions, we measured the level of the presynaptic protein synaptophysin. Synaptophysin is a good marker for synaptic integrity [41, 62, 63], and pathological synaptic alterations have been reported in PD post-mortem tissues [64]. Roughly 60% of PD patients suffer from cognitive impairments and dementia [65], indicating that their hippocampus, as a major region involved in memory formation, and their higher cortical association areas are affected. Finally, *in vitro*, addition to PFFs to cultured primary hippocampal neurons was reported to affect these neuron’s synaptic integrity and function [66]. Thus, we measured synaptophysin ipsi- and contralaterally in these brain regions in mice 90 dpi after PFF administration (Fig. 2). We found, in both regions, a highly significant, bilateral 20-25% reduction of this protein. Interestingly, we noticed this decrease in the absence of α-syn inclusions in the hippocampus. The α-syn oligomers though (Fig. 1) in that region may be linked to synaptophysin loss.

**Figure 2.**
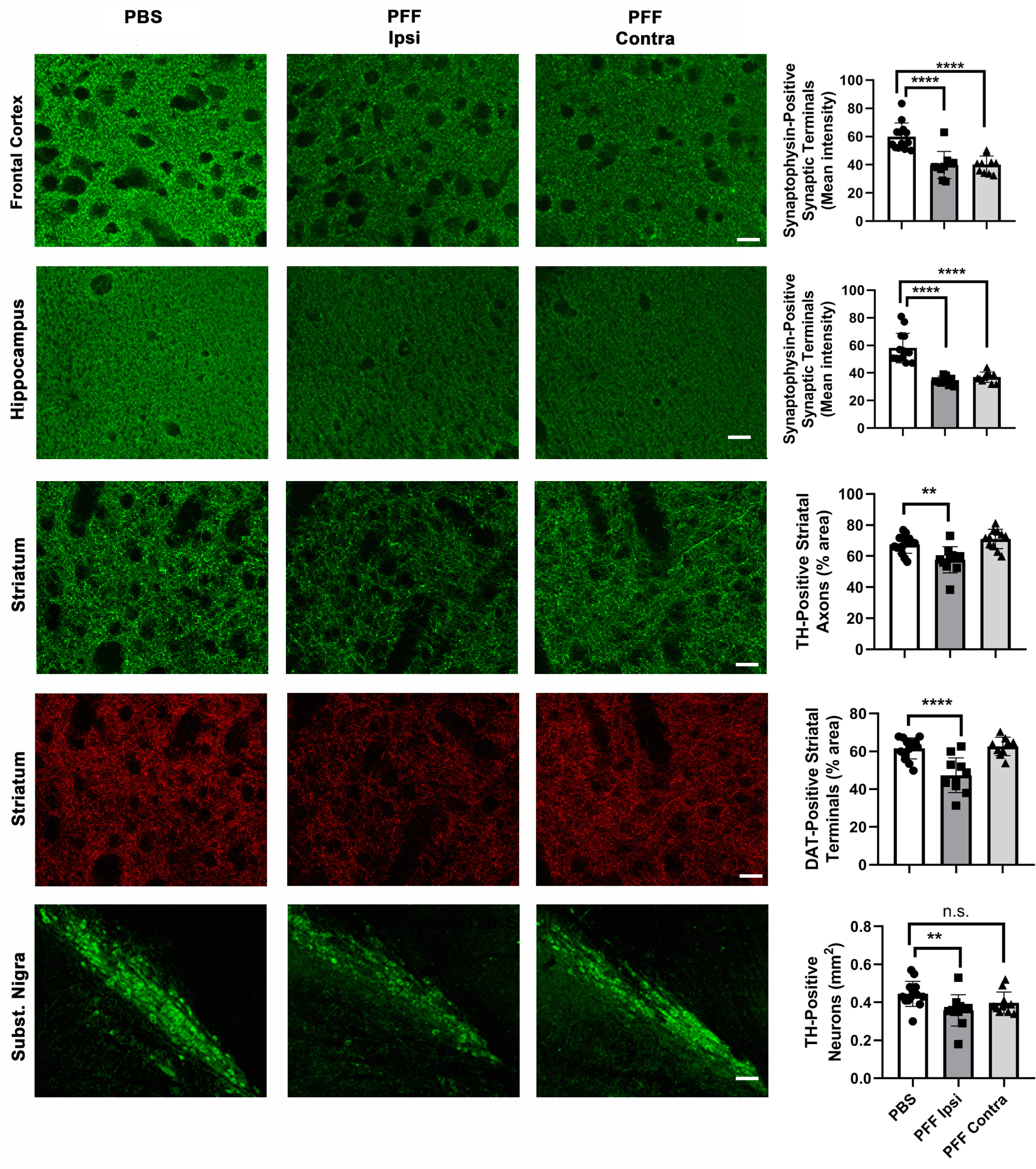
Intrastriatal injection of murine α-syn PFFs induced neurodegeneration in various brain regions. Mice were euthanized 90 dpi. In the frontal cortex and the hippocampus, a significant bilateral loss of synaptophysin-positive presynaptic terminals was observed (first two rows). In the striatum, a significant ispilateral loss of TH-positive axonal fibers and DAT-positive synaptic terminals was observed (3^rd^ and 4^th^ row). In the SN, a significant loss of TH-positive neurons was observed only ipsilaterally. For group comparisons and graphing, ipsilateral PBS measures were combined contralateral PBS measures, since they were similar. Pictures show the ipsilateral side of PBS-injected mice. **** p<0.0001, ** p<0.01, compared to PBS controls by Dunnett’s post-hoc; n = 10-11/group. Scale bars: 18 μm (for frontal cortical and hippocampal synaptophysin panels), 22.5 μm (for striatal TH and DAT panels), 80 μm (for Subst. Nigra panels)

Next, we examined the SN, because it contains dopaminergic neurons that are one of the most susceptible to PD-associated disease challenges. We measured the area occupied by tyrosine hydroxylase (TH)-positive neuronal profiles in the SN ipsilaterally, where α-syn inclusions were present (see above), but also contralaterally, which was without such inclusions. We did not find any sign of degeneration in the striatum or SN at 13 dpi (Supplemental fig. 2). We found though a 16% decrease of TH-positive neurons in the ipsilateral SN, that was significant, but not in the contralateral SN (Fig 2). To determine if striatal axonal projections of dopaminergic neurons were affected in our model, we analyzed the morphological integrity of these projections and their synaptic terminals. We observed, 90 dpi, a significant decrease in TH-positive axonal fibers as well as in dopamine transporter (DAT) positive synaptic terminals, in the ipsilateral, but not the contralateral striatum. To confirm ipsilateral striatal injury, we measured the neurotransmitter dopamine (DA) in dissected ipsi- and contralateral striata of PFF-injected and PBS control mice (n=8-12/group). We found a significant decrease in ipsilateral striatum of PFF mice compared their ipsilateral PBS controls (19.5+/−5.8 versus 27.5+/−7.3 pmol/mg; p=0.02 by ANOVA followed by Sidak’s posthoc, results are means +/− S.D.), but no difference between contralateral striatum of PFF mice compared to their compared their ipsilateral PBS controls (27.3+/−3.7 pmol/mg versus 28.5+/−5.7 pmol/mg). This conformed our histological observations.

### Widespread, pronounced, and bilateral microgliosis, caused by intrastriatal injection of α-syn fibrils

Microglia, the local CNS innate immune defense cells [67], react rapidly to CNS infection or injury in an orchestrated fashion. Functional imbalances of these cells can precipitate disease outcomes [36, 37, 68]. While strong microgliosis has been reported in PD and models thereof [69–71], the role of these cells in disease initiation and progression of PD is poorly understood.

To better understand the role of these cells in the context of α-syn spreading, and more precisely to determine if they have a role in driving the neurodegeneration we observed, we first analysed their response using a specific marker (Iba1), in mouse brains after injection of α-syn PFFs. We observed a surprisingly strong (4-5 times over control in some brain regions), microgliosis in brain regions with (bilaterally in frontal cortex, amygdala, SN) at 90 dpi. The microgliosis was present in brain regions with inclusions, but also those without (hippocampus) or very little (contralateral SN) inclusions (Fig. 3). While no significant Iba1 increase was seen at 90 dpi in the ipsilateral striatum in PFF injected mice, microglial Cluster-of-Differentiation 68 (CD68), a marker for phagocytic activity, was increased. No significant microgliosis was observed at 13 dpi (Supplemental fig. 2). Microglia in PFF-injected mice had thickened, though still ramified, processes, and an intensely stained cell soma. In the cerebellum, which was devoid of α-syn deposits in all mice, we could not detect any differences in Iba1 positive microglia between PFF-injected and control PBS injected mice (not shown).

**Figure 3.**
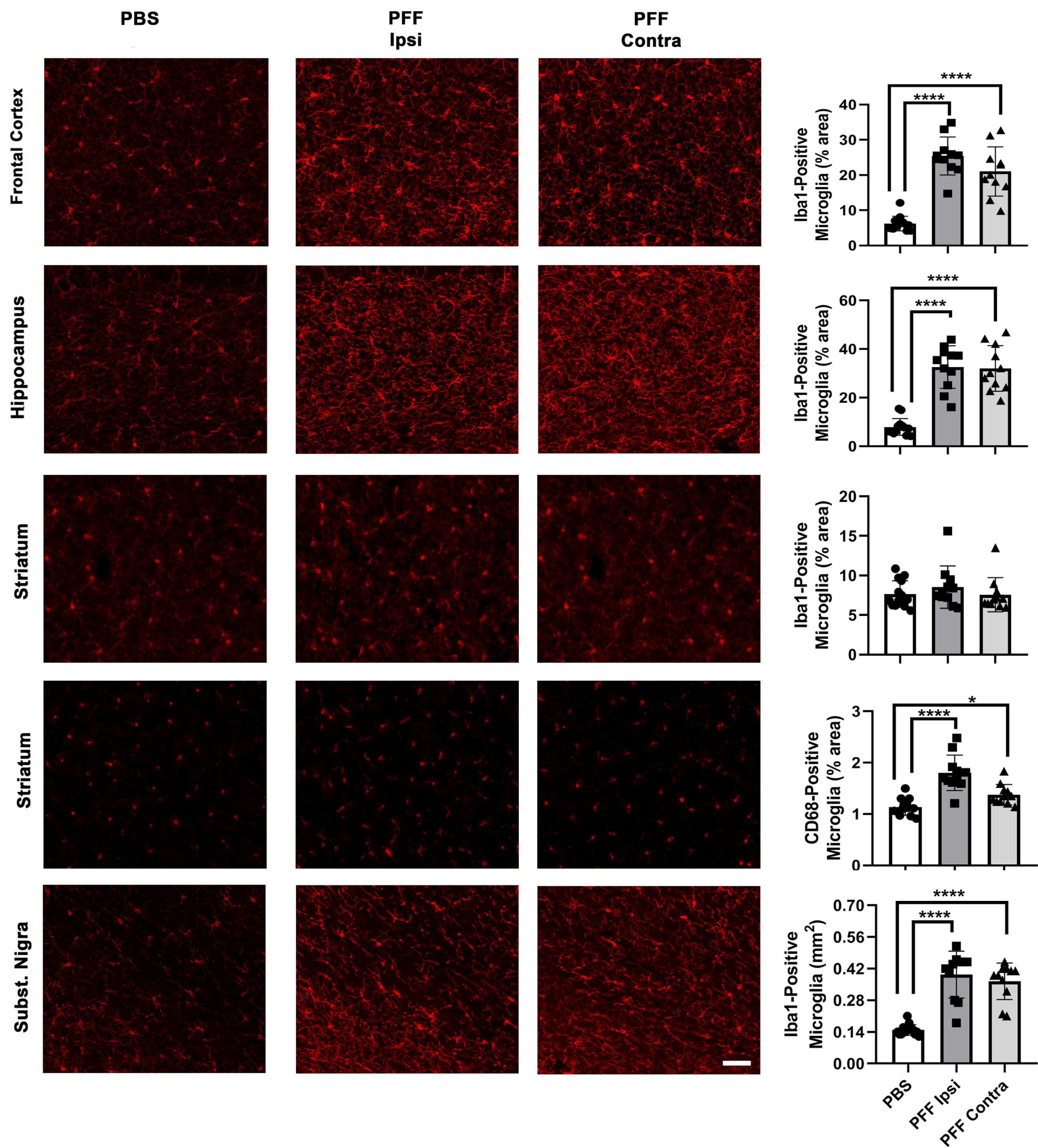
Intrastriatal injection of murine α-syn PFFs induced widespread microgliosis in different brain regions. Mice were euthanized 90 dpi. Panels show microgliosis measured on Iba1-stained sections of frontal cortex (upper row), hippocampus (second row), striatum (3^rd^ row), and Subst. Nigra (last row), and on CD68-stained sections of striatum (4^th^ row). A very strong microgliosis (up to 4x over control) was observed bilaterally in frontal cortex, hippocampus, and SN. No increase in Iba1 signal, but a significant bilateral increase in CD68 signal was observed in the striatum of PFF-injected mice. For group comparisons and graphing, ipsilateral PBS measures were combined contralateral PBS measures, since they were similar. Pictures show the ipsilateral side of PBS-injected mice. **** p<0.0001, * p<0.05, compared to PBS controls by Dunnett’s post-hoc; n = 10-11/group. Scale bars: 22.5 μm (for all panels).

Our observations indicate that a robust, widespread microglial reaction is an important part of the α-syn spreading process, and warranted further investigation into the pathological implications of that reaction.

### Neurodegeneration and microgliosis neither correlated with **α**-syn deposition, nor with each other

To gain insight into the pathological properties of α-syn inclusions, we correlated the inclusion load with neurodegeneration and with microgliosis measured locally in frontal cortex and SN. We found that inclusion load correlated with neither of the two (Fig. 4). Thus, neurodegeneration as well as microgliosis induced by α-syn PFFs were independent of α-syn deposition.

**Figure 4.**
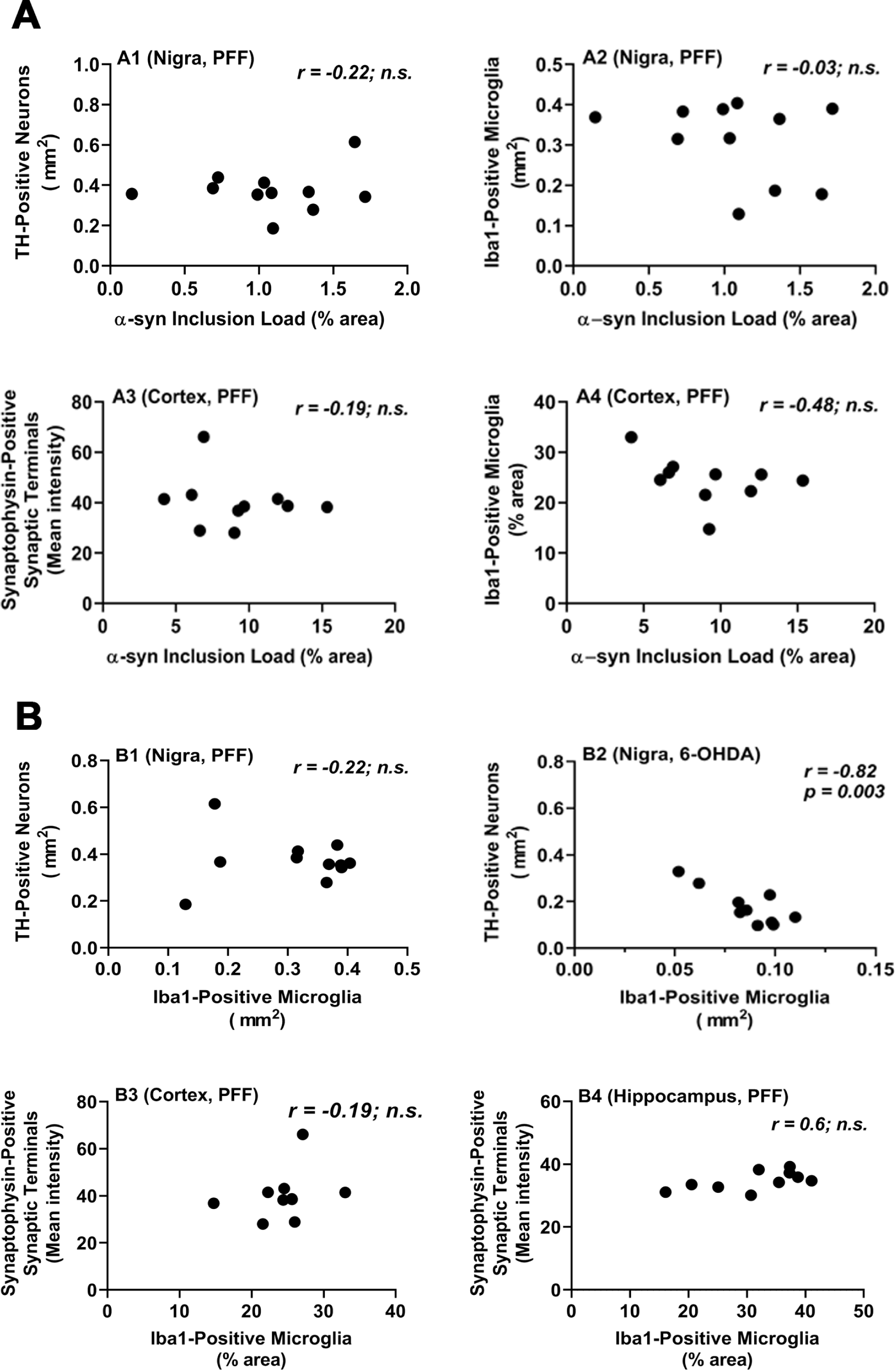
Different PD-related pathologies in the brains of mice injected intrastriatally with α-syn PFFs do not correlate with each other. Mice were euthanized 90 dpi. A. α-syn inclusion load did no correlate with neurodegeneration (loss TH-positive neurons, A1) or with microgliosis (A2) in the SN (Nigra), nor with neurodegeneration (loss of synaptophysin-positive synaptic terminals, A3) or with microgliosis (A4) in the frontal cortex (Cortex). B. Microgliosis did not correlate with loss of TH-positive neurons in the SN after intrastriatal PFF injection, but did so after intrastriatal injection of the toxin 6-OHDA. The microgliosis, measured on Iba1-stained section, was also much higher in the Subst. Nigra of PFF-injected mice than in that of 6-OHDA-injected mice. All measures shown are from the ipsilateral brain sides; similar observations were made for the contralateral sides of PFF-injected mice. Correlation analyses were done using Spearman rank test for data set including α-syn inclusion load measures, and with Pearson’s test for data sets with the other measures.

The strong microgliosis in different brain regions after administration of PFFs prompted us to look into this observation further. In the brain, microglia react rapidly to tissue injury to control the damage and clear up cell debris [37, 72]. Thus, microglial reaction is typically secondary to an underlying neurodegenerative process, and, as a consequence, increase of microglial reaction is directly associated with increase in neuronal damage. For instance, we have observed that microglial reaction (measured on Iba1 immunostained sections) correlated with TH neuron loss in the ipsilateral SN after unilateral 6-OHDA lesioning (Fig. 4B2), and with synapse or dendritic loss in the cortex after lesioning with the excitotoxin kainic acid [43]. After intracerebral injection of α-syn PFF though, we found that microglial reaction was not only much stronger than after injection of neurotoxins (4-5x *versus* 2-3x over control), but also failed to correlate with measures of neurodegeneration (TH neuron loss in the SN, synaptophysin in the cortex and hippocampus) (Fig. 4B). This observation indicates that the microglial reaction to α-syn spreading may be a direct response to factors produced during that process, and not just a secondary response to neuronal degeneration, as is the case after injection of neurotoxins.

### Microglia across several brain regions react strongly to intrastriatal injection of **α**-syn oligomers

Several studies have indicated that microglia are activated *in vitro* by α-syn oligomers [73, 74]. As described above, we have observed the presence of α-syn oligomers, notably in the hippocampus, after intrastriatal injection of α-syn PFFs. To test whether α-syn oligomers could be the factor that leads to a strong microglial reaction during the α-syn spreading process, we injected such oligomers into the same location as the PFFs, the dorsal striatum. Just 13 dpi, we observed, on Iba1 stained sections, a strong microglial reaction in the ipsilateral striatum, frontal cortex, and hippocampus (Fig. 5). Qualitatively, the reaction was even stronger than 90 dpi after PFF injection. This observation shows that, in a mouse brain, α-syn oligomers can induce a microglial reaction, even at a distance from the injection site. Thus, α-syn oligomers emerge as the likely factor that, by diffusing through the brain, induces a strong microglial reaction during the process of α-syn spreading.

**Figure 5:**
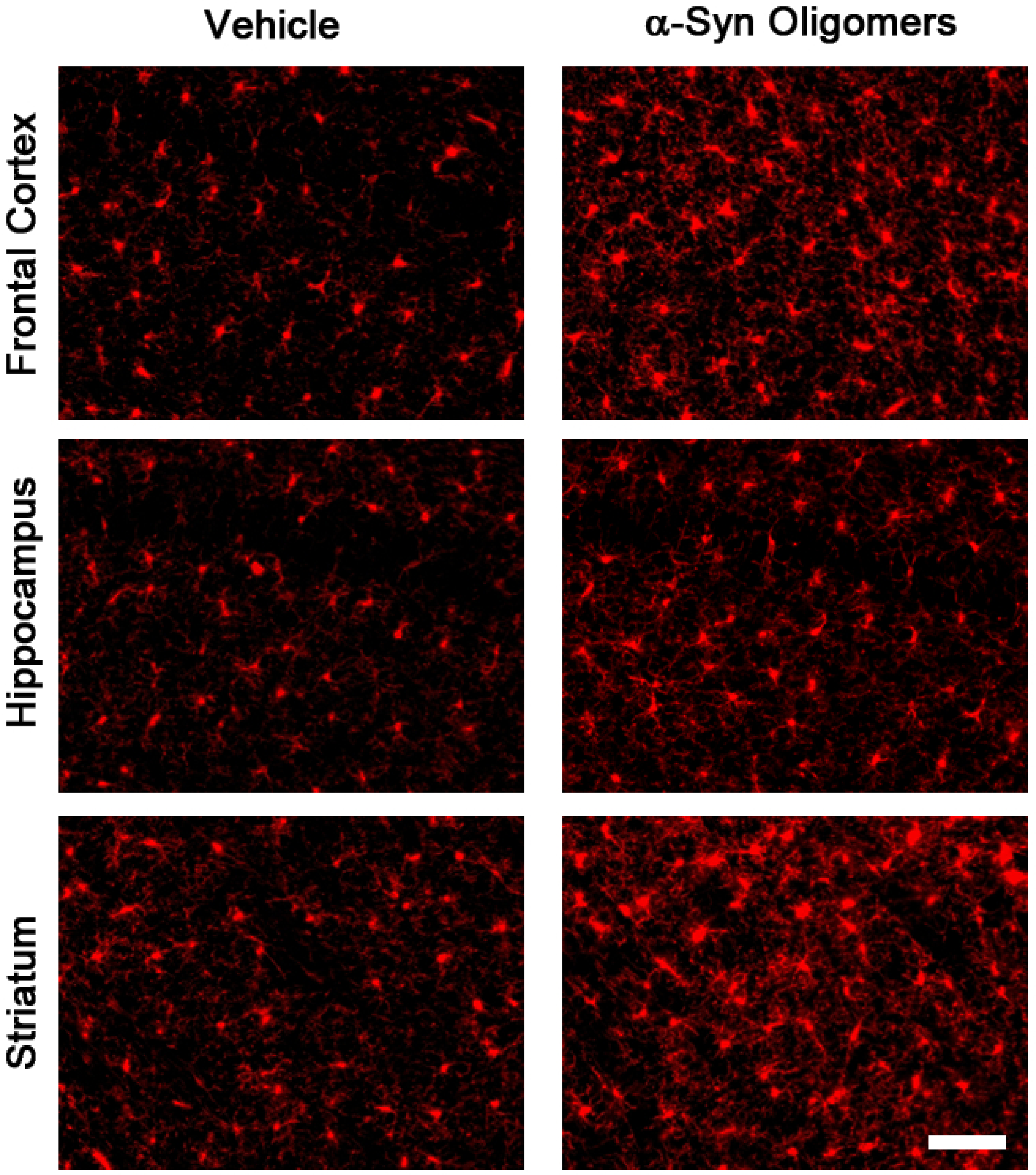
Strong microglial response after intrastriatal injection of α-syn oligomers. Oligomers were prepared and injections were performed as described in Materials and Methods. A strong microgliosis was observed in different brain regions 13 dpi, showing that α-syn oligomers are robust microglial activators in vivo. Scale bar = 40 μm.

### Transcriptional profiling of ventral midbrain reveals most gene expression changes occur 13 days after **α**-syn PFF injection

To investigate the molecular underpinnings of the neurodegeneration and of the microglial response accompanying α-syn spreading, we generated a gene expression profile from ventral midbrain of PFF injected and control mice using the Affymetrix gene expression profiling platform. Because microglial response typically starts early after an insult [67, 75], we analyzed the midbrain gene expression profiles 13 dpi (no neurodegeneration) and 90 dpi (neurodegeneration in the ipsilateral striatum and midbrain) after intrastriatal α-syn PFF injection.

We focused on two comparisons of ventral midbrain gene expression profiles: 1. ipsilateral midbrain of PFF-injected mice (ipsi PFF, with degeneration of nigral TH neurons and their striatal projections) *versus* ipsilateral midbrain of control PBS-injected mice (ipsi PBS), 2. ipsilateral midbrain of PFF-injected mice *versus* contralateral midbrain of the same, PFF-injected, mice (contra PFF, without loss of nigral TH neurons and their striatal projections). We figured that these two comparisons would be best suited to reveal relevant gene expression changes.

The number of DEGs that emerged in these comparisons, and the number of overlapping DEGs between the two time points (13 dpi and 90 dpi) are shown in Fig. 6. By comparing ipsi PFF to ipsi PBS, applying a cut-off of p<0.05, we found a total of 2.631 significant DEG at 13 dpi, and significant 2584 DEG at 90 dpi, with 985 overlapping DEGs between the two time points. After correcting for multiple hypothesis testing at a cut-off of pfp < 0.1, we found 266 DEGs at 13 dpi, and 82 DEGs at 90 dpi, with 39 DEGs overlapping between the two times points. The majority of overlapping DEGs showed upregulation at 13 dpi, but downregulation at 90 dpi (Venn diagrams of overlapping DEGs in Fig. 7), indicating active gene expression at the early time point after PFF injection. By comparing ipsi PFF to contra PFF, we found 3.477 significantly DEGs at 13 dpi, and 3.209 DEGs at 90 dpi, with 1356 overlapping DEGs between the two time points. At pfp < 0.1, we found 648 DEGs at 13 dpi, and 588 DEGs at 90 dpi, with 227 overlapping DEGs. At 13 dpi, we found a similar number of upregulated *versus* downregulated DEGs, but at 90 dpi, we saw that most DEGs were, interestingly, downregulated.

**Figure 6.**
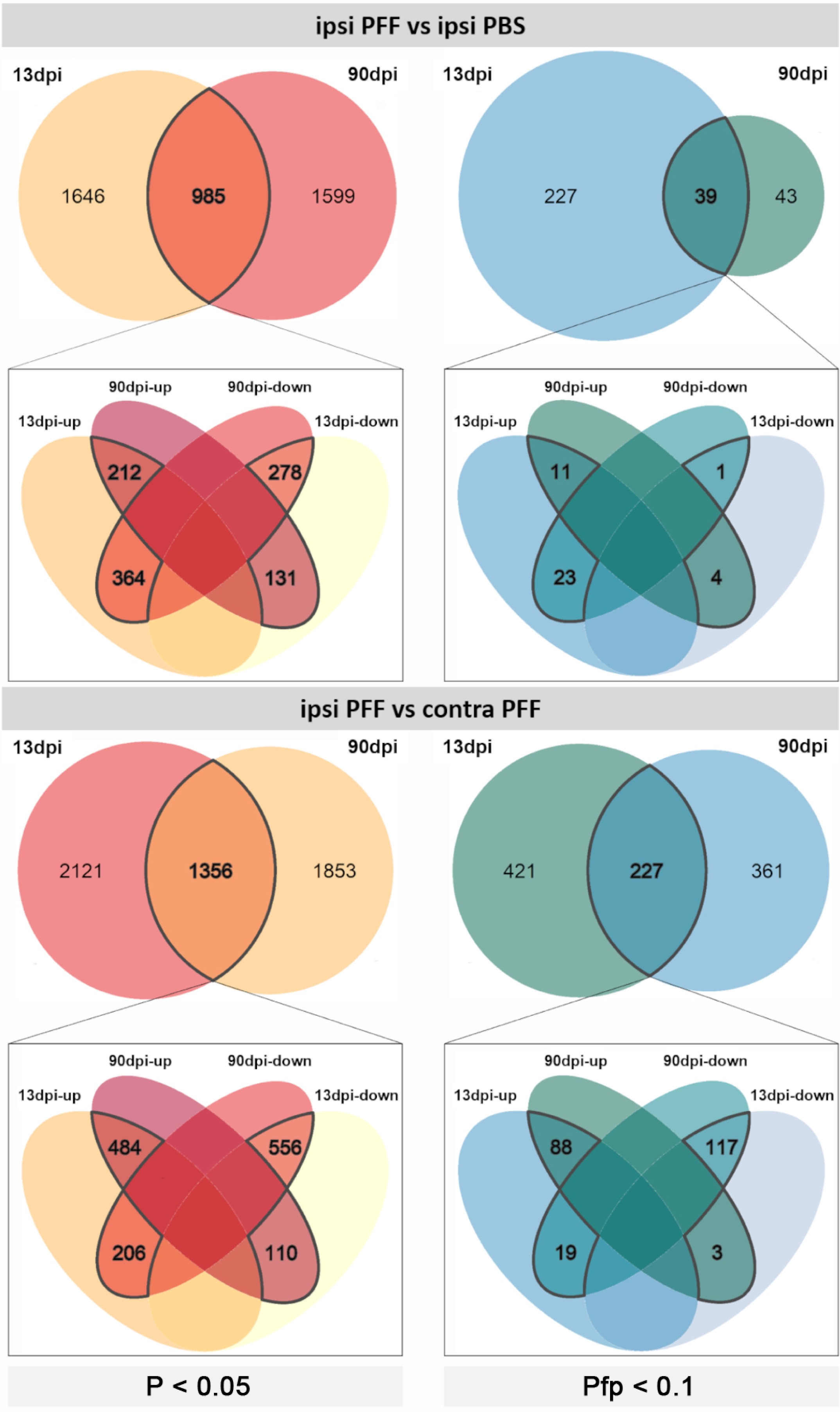
Differentially expressed genes induced in the ipsilateral ventral midbrain of mice at 13 dpi (A) and 90 dpi (B) after intrastriatal injections of α-syn PFFs. Comparisons for each time point were made between gene expressed in ipsilateral ventral midbrain of PFF-injected mice versus those in the ipsilateral ventral midbrain for PBS-injected mice (ipsi PFF *vs*. ipsi PBS, n =6/group; upper 4 panels), and between gene expressed in ipsilateral ventral midbrain *versus* the contralateral ventral midbrains of PFF-injected mice (ipsi PFF *vs.* contra PFF, n =6/group; lower 4 panels). Panels on the left show Venn diagrams with numbers of DEGs with a significance level of p<0.05 (n=6 mice/group). Because some gene products have an effect within biological pathways while the expression of their genes may only change minimally, we show DEGs that emerge with this level of statistical stringency. In ipsi PFF *vs.* ipsi PBS, the number of DEGs was 2631 at 13 DPI, and 2584 at 90 DPI, with 985 DEGs common to both time points. In ipsi PFF *vs.* contra PFF, the number of DEGs was 3477 at 13 DPI, and 3209 at 90 DPI, with 1356 DEGs common to both time points. Panels on the right show Venn diagrams with the number of DEGs after adjusting for multiple hypothesis testing at pfp<0.1. In ipsi PFF *vs.* ipsi PBS, the number of DEGs was 266 at 13 DPI, and 82 at 90 DPI, with 39 DEGs common to both time points. In ipsi PFF *vs.* contra PFF, the number of DEGs was 648 at 13 DPI, and 588 at 90 DPI, with 227 DEGs common to both time points. The Venn diagrams below the main ones indicate the number of DEGs that show enhanced (“up”) versus decreased (“down”), as well as the overlaps, in the different comparisons.

**Figure 7.**
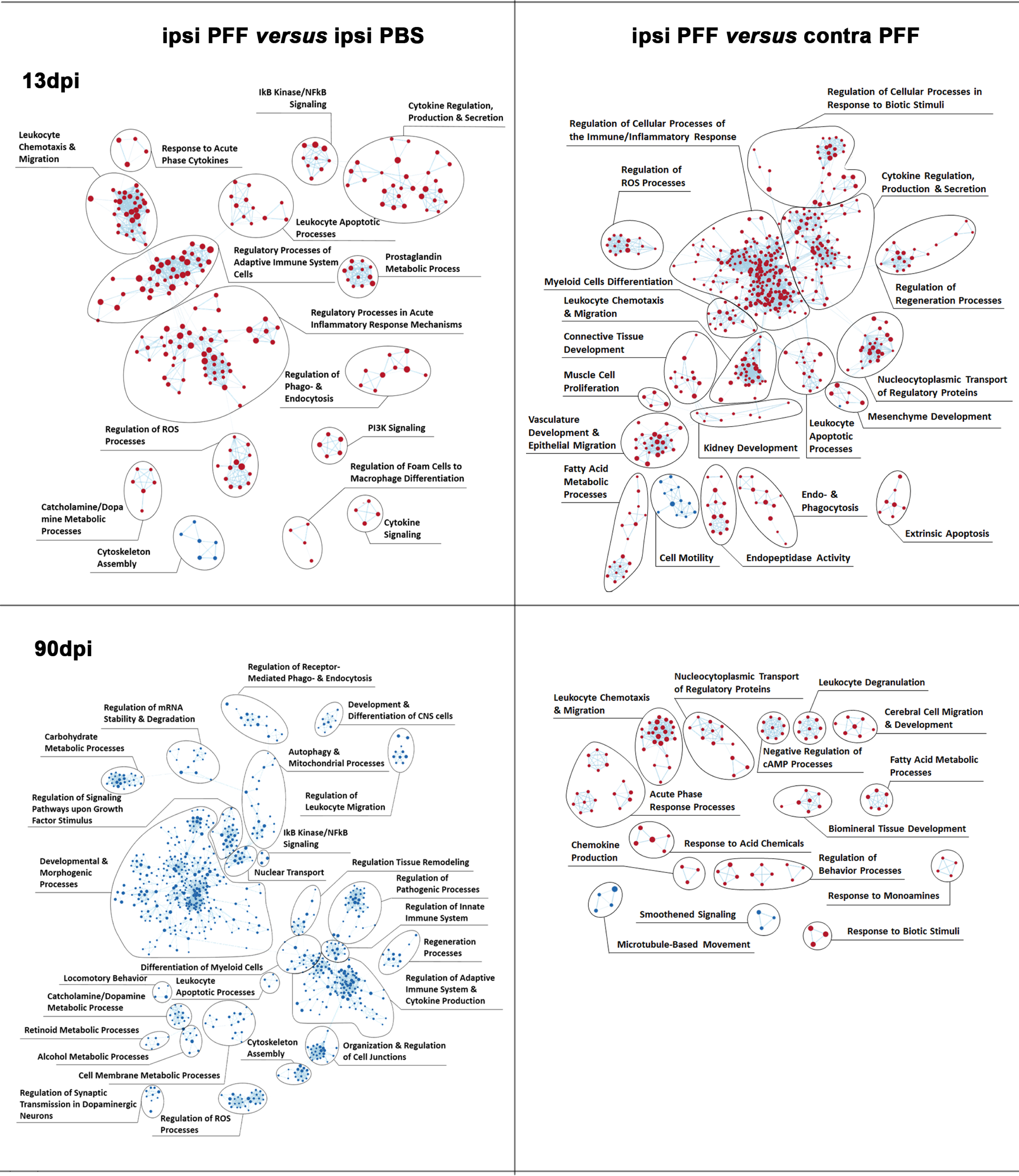
Enriched inflammatory pathways precedes neurodegeneration in mouse ventral midbrains after intrastriatal α-syn PFFs injection. Enrichment map of gene expression profiles were derived from GSEA. Statistics were done by weighted Kolmogorov-Smirnov, gene set size limits were set to min15 – max250. Details of curation procedure used to group BPs (represented as dots, either red if upregulated, or blue if downregulated) into high-level functional gene set clusters of BPs of related biological function are described in Material & Methods. At 13 dpi, comparing ipsi PFF to either ipsi PBS or contra PFF, most BPs were upregulated and associated with gene sets related to immune and inflammation processes. This shows that, in the ipsilateral nigro-striatum, neuroinflammation precedes neurodegeneration (measurable at 90 dpi), and might contribute to its development. At 90 dpi, comparing ipsi PFF to ipsi PBS, all BPs, including those associated with inflammation gene sets, were downregulated, possibly reflecting the neurodegenerative process itself. Comparing ipsi PFF to contra PFF at this time point, most BPs were upregulated.

Taken together, these two comparisons indicate that enhanced gene expression changes occurred in the ventral midbrains of both hemispheres at 13 dpi, probably setting the stage for the subsequent pathological events. In contrast, at 90 dpi, in the ipsilateral midbrain, most DEGs dial their expression level back, indicating a reduction in gene transcription, while the major pathological events now appear to take place at the protein level, and are measurable with quantitative histology (see above).

### Gene set enrichment revealed early involvement of inflammation in the **α**-syn seeding/spreading process

To investigate which molecular pathways underlie the α-syn spreading process and its associated pathologies, in particular microgliosis, we generated an enrichment map, using Gene-Set Enrichment Analysis (GSEA, see Methods), with each gene set based on a Biological Process (BP, see Methods for details). To obtain a global view of the BPs alterations during the evolution of α-syn spreading induced pathologies, we used manual curation to group gene sets into biologically meaningful gene clusters associated with high order pathological processes (Fig. 7).

Our first observation was that, in ipsilateral midbrains of PFF-injected mice compared to those of PBS-injected ones, 261 BPs were enriched at 13 dpi, but, surprisingly none at 90 dpi. In contrast, we observed that, at 90 dpi, all BPs in ipsilateral midbrains of PFF-injected mice *versus* those of PBS-injected ones (total of 1067 BPs), showed reduced gene activity. This observation indicates a significant shift from enhanced to greatly reduced transcriptional activity in the time frame between 13 dpi and 90 dpi, and confirms the observations on DEGs depicted in Fig. 6.

We then observed that many gene clusters with enhanced transcriptional activity at 13 dpi in ipsi PFF were associated with inflammation/immune processes (Fig. 7, upper panels), while gene clusters associated with similar activities had reduced transcriptional activity at 90 dpi, in particular compared to ipsi PBS (Fig. 7, lower left panel). This indicated that, after an initially enhanced activity of genes regulating inflammation/immune responses, that activity was strongly reduced at a stage when pathology was detectable histologically.

Another interesting observation we made was that some gene clusters containing BPs associated with reduced gene activity at 90 dpi, were related to dopaminergic neuron activity (e.g. catecholamine/dopamine metabolic processes, locomotor behavior, regulation of synaptic transmission regulation of signaling pathways upon growth factor stimulus). The reduced gene activity in midbrain dopaminergic neurons was likely a reflection of their pathological demise.

Taken together, these observations point to an important role for inflammatory/immune processes, in the initiation and progression of neurodegeneration in the context of α-syn spreading.

### Gene expression changes confirm early microglial reaction in response to **α**-syn seeding/spreading

To identify the immune cell type(s) active in the inflammatory response to α-syn seeding/spreading, we looked at the 20 most highly changed DEGs and their cellular source for each time points after PFF injection (Fig. 8). We used a public database based on single cell expression profiling from mouse brain to assign a cell type to each DEG in our list (see Methods).

**Figure 8.**
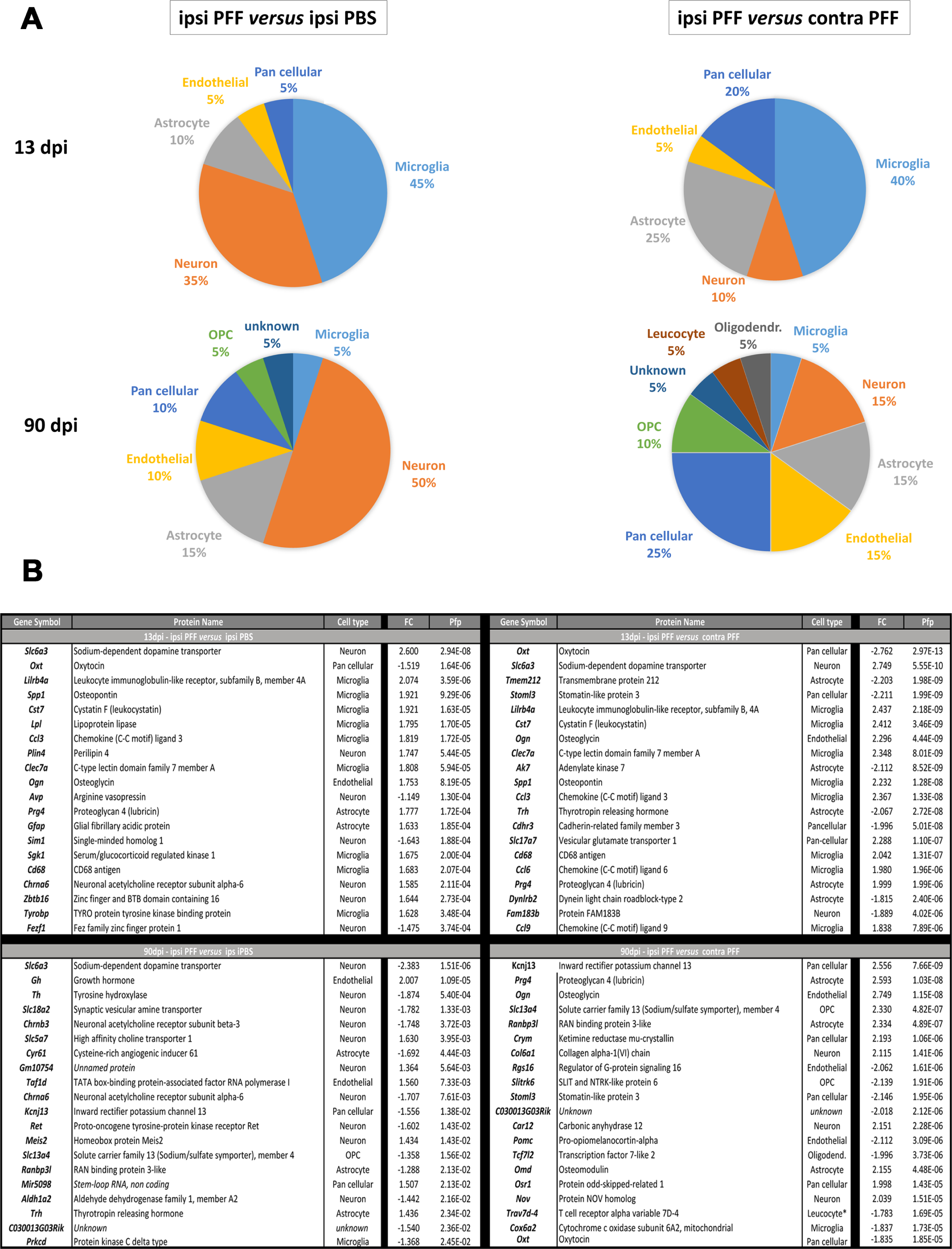
Top 20 DEGs in mouse ventral midbrain after intrastriatal injection of α-syn PFF indicate involvement of microglia in initial pathological events. Cellular source of DEGs was determined using the brain RNAseq database: https://www.brainrnaseq.org/. The top panels show pie charts with the cellular source of the 20 top DEGs when comparing ipsi PFF to ipsi PBS (left panels) or ipsi PFF with contra PFF (right panels) at 13 dpi and 90 dpi. At 13 dpi, comparing ipsi PFF with ipsi PBS or contra PFF, 45% and 40%, respectively, of the top 20 DEGs were microglial. At 90 dpi, comparing ipsi PFF with ipsi PBS, 50% of top 20 DEGs were neuronal, possibly a reflection of neurodegeneration. The bottom panel lists the gene products of the gene symbols, coded proteins, the associated cell type 20, the fold change (FC) and the pfp of all top 20 DEGs for each comparison.

At 13 dpi, in both the ipsi PFF *versus* ipsi PBS as well as the ipsi PFF *versus* contra PFF comparison, we observed that the majority of genes with enhanced expression were microglial (ipsi PFF *versus* ipsi PBS: 9 out of 20, or 45%, ipsi PFF *versus* contra PFF: 8 out of 20, or 40%). This indicates a strong gene expression activity of these cells, well before morphological changes can be detected histologically.

In contrast, at 90 dpi, we observed that only 1 out of 20 (5%) of DEGs was microglial in both comparisons (ipsi PFF *versus* ipsi PBS, ipsi PFF *versus* contra PFF). The majority (50%) of DEGs in the ipsi PFF *versus* ipsi PBS comparison were neuronal.

The observation that the majority of DEGs at 13 dpi were microglial, confirmed an early and strong response of these cells, at least on the molecular level.

### Unusual microglial molecular signature, induced by intrastriatal injection of **α**-syn PFFs, precedes neurodegeneration

To better understand the microglial molecular processes accompanying the α-syn spreading process, we looked at expression of a series of genes coding for factors associated with typical pro-inflammatory (M1) or anti-inflammatory (M2) profile [76], with the same comparison pairs as before: ventral midbrains ipsi PFF *versus* ipsi PBS, and ventral midbrains ipsi PFF *versus* contra PFF (Table 1). In our model, we observed no clear-cut pro-inflammatory M1 nor anti-inflammatory M2-profile. Interestingly, we also observed no evidence, at 13 dpi or 90 dpi, for gene expression changes in classical M1 markers such as *Il1b*, *Tnfa, or Nos2*. M1 markers whose gene expression was enhanced were *Cybb*, *Ptgs2*, and *Cxcl10*. NADPH oxidase 2 (Nox2), coded by *Cybb*, generates free oxygen radicals, which can harm neurons [77]. Cyclooxygenase 2 (Cox2), coded by *Ptgs2*, generates arachidonic acid metabolites, some of which have been reported to be neurotoxic [78] or form neurotoxic dopamine-quinone adducts [79]. Thus, this may be mediators of the neurodegeneration at 90 dpi observed in ipsi PFF midbrains. The only M2 marker that showed enhanced gene expression was *Mrc1*, coding for the mannose receptor. Other microglial activation markers that have not been associated specifically with an M1 or M2 profile though, such as *Cd68*, *Tyrobd*, *Trem2*, *Tlr2*, *P2ry6*, and *Aif1,* showed increased expression at 13 dpi and/or 90 dpi. *Mrc1*, *Cd68, P2ry6, Aif1* gene products are all involved in phagocytic processes and/or signal transduction [80–82]. We had observed CD68 upregulation by immunostaining in the striatal projection area of dopaminergic neurons (Fig. 3). The *Tlr2* gene product is a receptor for α-syn, an interaction that elicits the production of microglial neurotoxins [83], and anti-TLR2 antibody administration has been reported to have therapeutic efficacy in mouse models of α-syn toxicity [84]. *Tyrobp* and the gene for its receptor, *Trem2*, whose product is involved, among other processes in the regulation of microglial phagocytosis [85], were also upregulated.

Finally, to see if there was an astroglial and peripheral immune cell involvement in our model, we listed gene expression data for typical markers of these cells from our gene expression dataset (Supplemental table 3). Enhanced expression of a series of astroglial genes in ipsi PFF midbrain indicates a reaction of these cells. Enhanced expression of *Ptprc*, which codes for CD45, a marker that can be both expressed by microglia and invading macrophages, and of *Cd4*, which codes for the helper T cell antigen CD4, in the same region, indicates possible infiltration of peripheral immune cells that could contribute to neuronal injury [86–88]. Overall, we conclude that this unique molecular signature in the ipsilateral ventral midbrain at 13 dpi underlies the initial molecular events that lead to the neurodegeneration we observed at 90 dpi. Since neurodegeneration in the contralateral SN has been reported at later time points after PFF injection previously [20], in one study even in the absence of α-syn inclusions [89], it is reasonable to assume that, at a point past 13dpi, the same molecular signature occurs there.

Our data indicate that inflammatory events, in particular those associated with microglia, and not inclusion formation, are initiators of neurodegeneration in the context of α-syn spreading in PD.

## Discussion

In this study, we have used a seeding/spreading model of α-syn, based on striatal injection of PFFs in the mouse brain, to investigate key questions on how two major pathological features of PD, α-syn inclusion formation and neuroinflammation, contribute to neurodegeneration. We provide evidence that: 1. α-syn inclusion formation does not correlate with neurodegeneration: in areas with inclusions, the inclusion load did not correlate with the extent of neurodegeneration, and, in at least one area (hippocampus), neurodegeneration was detected in the absence of inclusions; 2. an exceptionally strong microglial response was seen across different brain regions, but this response did also not correlate with neurodegeneration; 3. the most likely driver of the microglial response were diffusible α-syn oligomers; 4. gene expression changes indicative of early neuroinflammatory events in the ventral midbrain, in particular in microglia, appeared before nigro-striatal degeneration, and some of these factors could be the driver for downstream neurodegeneration. Our study provides novel insights into underlying pathological processes of α-syn spreading mediated PD-like neuronal injury.

We undertook this study because it is unclear how different pathological processes relate to each other in PD. In particular, it is intensely debated whether the α-syn spreading and inclusion formation are the main driving forces in disease initiation and progression, or whether other processes do this or at least participate in them [90–92]. While the nature of the initial trigger of α-syn’s misfolding and seeding is still unknown, the hypothesis that its spreading and seeding in a “prion-like” fashion along interconnected neuronal pathways, ultimately leading to inclusions, is a major driver of the PD pathological process has gained momentum [8, 93]. Observations on PD patients have provided indirect evidence for that view. For instance, the Braak hypothesis [10] posits that α-syn pathology starts in lower motor nuclei of the brainstem (such as the Dorsal Motor Nucleus of the Vagus) or even in the PNS, then gradually moves upwards and, in doing so, causes various PD symptoms, from early non-motor to later motor and cognitive and psychiatric ones, to appear. Other routes of propagation, such as starting and spreading out from the olfactory bulb, have also been suggested [8]. Such a gradual progression of disease could elegantly be explained by α-syn spreading like a prion. It was also observed that fetal grafts of dopaminergic neurons into the striatum of PD patients develop α-syn inclusions after a few decades, which they could have acquired as consequence of α-syn spreading from the surrounding disease tissue [12]. More direct evidence for the importance of α-syn spreading in inducing PD-like disease comes from experimental models. In rodent or primate models, direct injection, in different brain regions, of PD brain tissue, isolated Lewy bodies, or PFF made out of recombinant α-syn, or viral vector driven local overexpression of α-syn, induces a variety of PD-related pathologies, including α-syn spreading along connected neurons and inclusion formation [19]. Peripheral PFF injections, such as intramuscular or intestinal, have been reported to also lead to PD-like pathologies in the brain of mice [94–96]. These studies have cemented, experimentally, the process of “prion-like” propagation and inclusion formation of α-syn.

The mechanism of this process has been investigated in *in vitro* systems. Cultured neurons secrete as well as take up circulating α-syn, and various underlying mechanisms have been proposed, such as unusual forms of endo- and exocytosis, or nano-tubes [97]. Ingested, presumably misfolded α-syn, corrupts its endogenous counterpart and leads it to form pathological inclusions [58]. During the process, different neuronal functions, such as axonal transport or mitochondrial respiration, get impaired, neurons malfunction and may ultimately die [98, 99]. Glial cells have also been reported to take up α-syn, and, in some cases, this can lead to pathological inclusions as well as in the case for oligodendrocytes in Multiple System Atrophy [100]. The toxic potential of inclusions has also been investigated *in vivo*. In a mouse model of α-syn spreading, where inclusion formation was followed *in vivo* by multi-photon laser microscopy, the formation of intraneuronal inclusion was reported to coincide with neuronal dysfunction [33]. Another study has shown a weak correlation between loss of TH neurons in the SN and a global score of inclusion load after striatal PFF injection in both mice and rats, but a strong correlation between the two measures after direct injection of PFFs into the SN of rats [34]. Thus, it is tempting to conclude that α-syn inclusions are at the very least one major driver of PD pathologies. But a closer look at other evidence reveals several unresolved questions in this otherwise elegant picture. In post-mortem brain tissues of early or late PD, the correlation between α-syn inclusion (Lewy body) load and nigral degeneration is not always clear [25, 26]. Across different studies looking at various brain structures affected in PD, α-syn inclusions have been reported in areas with high, moderate, or no neuronal loss [26]. Some PD patients, including some familial forms, have PD symptoms and loss of nigral neurons without detectable α-syn inclusions [27]. Some neurons, such as GABAergic neurons, while appearing in the path of α-syn spreading, never develop inclusions [27]. Interestingly, one study, comparing Incipient Lewy Body Disease (ILBD) to PD autopsy material, reported that neuronal loss precedes α-syn inclusion formation in the SN [101]. In a rodent model where spreading is driven by viral overexpression of α-syn in the Dorsal Motor Nucleus of the Vagus, while intact neuronal architecture was essential for the spreading process to happen, neurodegeneration and inclusion formation were also found to be independent processes [61].

Non-fibrillar forms of misfolded α-syn, notably oligomers, diffusing for long distances within the brain, have been suggested to drive neuronal dysfunction and degeneration [30, 90]. In our study, we indeed found evidence for neurodegeneration that was independent of the presence of inclusions, and, in the hippocampus, even appeared in the complete absence of those, but in the presence of oligomers. Published evidence suggests that the hippocampus remains devoid of α-syn inclusions even 180 days after PFF injection into the striatum [20]. Our data therefore does not support the notion of a direct relationship between the formation of α-syn inclusions and neurodegeneration, but rather indicate that the α-syn spreading process may lead to the formation of pathological oligomers that may be the driver of PD-like neurodegeneration.

Pathologically misfolded α-syn can drive neuronal injury in PD by different means, and those include mitochondrial dysfunction, oxidative stress, endoplasmic reticulum stress and lysosomal dysfunction, disequilibrium in cytosolic Ca^2+^, neurotoxic oxidized dopamine, disruption of axonal transport, and neuroinflammation [99]. The relative contribution of these different processes to neuronal demise is unclear. Neuroinflammation though has received particular attention because of its widespread involvement in various neurological diseases and the potential for therapeutic modulation [75, 102, 103]. The major cellular mediators of this process are microglia. Microglia are a particular kind of myeloid cells that originate from the yolk sack and populate the nervous system during early stages of development, where they act as the innate, resident immune cells [67, 82]. During development and under normal conditions, they modulate nervous system homeostasis, prune synapses and regulate their formation. Under pathological conditions, they act as the primary line of defense against infectious organisms, and clear endogenous tissue debris after injury [37, 67, 82]. They undergo a substantial morphological and functional transition to activated, or reactive, microglia, which makes them functionally equivalent to macrophages [67]. Evidence suggests though that, in many neurological conditions, they are not only reacting to disease, but play an active part in tissue injury exacerbation and propagation [82]. This pathological process is, in particular in PD, incompletely understood. While microglial activation can be induced by neuronal injury and/or misfolded and aggregated protein, notably α-syn oligomers or fibrils [104], it is still unclear how and when microglial activation damages healthy tissue and exacerbates the neurological disease process. In PD, a strong microgliosis is observed *post mortem* in the SN [70, 105]. Longitudinal imaging studies with PET ligands demonstrated an early microglial activation in various regions beyond the SN, such as cortex, hippocampus, basal ganglia, and pons, but no correlation with other pathological measures, including clinical scores, of PD emerged [106, 107]. Interestingly, in striatal fetal grafts implanted in PD patients [12], microglia activation was observed years before the appearance of α-syn inclusions [108]. In different toxin-induced PD rodent models, microgliosis was reported to precede, coincide, or follow the appearance of neuronal demise[70], while in a transgenic human α-syn model [109] and in rats injected with PFFs into the striatum [110], microgliosis, measured histologically, was shown to precede neurodegeneration.

These studies are based on the observation mainly, if not exclusively, of morphological changes of microglial response using immunostaining techniques for generic cell markers. While informative, the detection of morphological changes indicating microglial activation does not yield enough information on the actual physiological or molecular profile of these cells. The common distinction to characterize two functional states of activated microglia is the M1/M2 terminology, with M1 representing a pro-, whereas the M2 representing an anti-inflammatory activation status [76]. This distinction however is often inadequate as microglia commonly have a spectrum of activation states that may change over the course of the disease [111]. Recent gene expression profiling approaches have revealed a bewildering complexity in microglial heterogeneity [36, 112]. Evidence suggests a “core” gene expression profile response that is associated with every neurodegeneration condition, while expression changes of a more restricted set of genes may be specific for each condition, leading to the concept of disease-specific microglial signatures [112]. Our study provides new insights into the molecular underpinnings of neuroinflammation preceding neuronal injury in a PD-like context of α-syn spreading, and points to an active role of microglia in inducing neurodegeneration. First, we show, at the level of gene expression, that neuroinflammation-linked processes are activated, and that many microglial genes had increased expression levels at an early (13 dpi), which then were down-regulated at a later (90 dpi) time point after PFF injection. Microglia genes that code for factors causing neurodegeneration showed increased expression 13 dpi only in the ipsilateral midbrain, where TH loss was observed later, at 90 dpi. Among these were *Cybb,* which codes for NAPDH oxidase 2, an enzyme that catalyses the production of tissue harming free radicals [77], and *Ptgs2,* which codes for cyclooxygenase 2 (Cox2), an enzyme that forms prostanoids from arachidonic acid, some of which are neurotoxic [78, 113]. Interestingly, *Tlr2*, *Trem2*, and *Tyrobp* RNAs showed increased levels in our model at 13 dpi. Many genes linked to microglial activation are regulated by Tyrobp, a tyrosine kinase binding protein acting that binds to Trem2. The Tyrobp/Trem2 pair triggers pathways that are involved in the inhibition of TLR-mediated inflammation, and in modulating phagocytosis [85]. *Tlr2*-deficient mice are protected against neurodegeneration induced by transgenic α-syn overexpression [114]. In prodromal PD, TLR2 immunoreactivity on microglia was reported to be enhanced, whereas in late stage PD, it wasn’t [115], indicating that, just like in our model, the microglial response happens in early phases of the disease and changes over time. Alpha-syn, in particular in its oligomeric form, activates microglia *in vitro* through Toll-like receptors [73, 104], and targeting TLR2 by immunotherapy was shown to be beneficial in α-syn pathology models [84].

The absence of increased gene expression of common pro-inflammatory mediators such as *Il1b* and *Tnfa* in our α-syn spreading model is puzzling, since these are factors associated with most, if not all, inflammatory conditions. Of note though is that we also did not observe enhanced expression of these factors when primary microglia where exposed to our α-syn PFFs, while they responded strongly to bacterial lipopolysaccharide (not shown). It is possible that the increased expression for these genes was missed and occurs at a time point after PFF injection that we haven’t looked at, or that they are indeed not expressed in this model. Overall, the gene expression signature of microglia we detected was neither typical pro-inflammatory M1-nor anti-inflammatory M2-like, and our findings give further credence to the notion that microglia evolve on a spectrum of functional states as the disease progresses.

Taken together, our data indicate that, at least in the initial period of PD-like disease progression that is associated with α-syn spreading, non-deposited pathological forms of α-syn, such as oligomers, drive neurodegeneration in different brain regions *via* their action on microglia. Triggered microglia respond early, before neurodegeneration is apparent, by producing neurotoxic compounds, and through what appears to be a series of different activation states as the disease progresses. Our findings contribute toward first answers to key unresolved questions around neuroinflammation in PD [102], and have important implications for the design of therapeutic interventions during the early stages of the disease.

## Abbreviations

6-OHDA: 6-Hydroxydopamine
13 dpi: 13 days post injection
90 dpi: 90 days post injection
Aβ: amyloid beta peptide
AD: Alzheimer’s disease
α-syn: alpha-synuclein
BP: biological process
CD68: cluster of differentiation 68
Contra: contralateral
COX2: Cyclooxygenase 2
CNS: central nervous system
DA: dopamine
DAT: dopamine transporter
DEG: differentially expressed gene
ES: enrichment score
FDR: false discovery rate
GO: gene ontology
GSEA: gene set enrichment analysis
Iba1: ionized calcium binding adaptor molecule
Ipsi: ipsilateral
PBS: phosphate-buffered saline
PET assay: paraffin-embedded tissue assay
PFF: Pre-formed fibril
Pfp: percent false positives
PLA: proximity ligation assay
p-SER129-α-syn: alpha-synuclein phosphorylated at serine position 129
SN: Substantia Nigra
TDP43: TAR DNA-binding protein 43
TH: tyrosine-hydroxylase
TLR2: Toll-like receptor 2
WB: Western blot

## Additional information

### Ethics approval

Animal studies performed at the Luxembourg Centre for Systems Biomedicine were approved by the institutional Animal Experimentation Ethics Committee of the University of Luxembourg, and the responsible Luxembourg government authorities (Ministry of Health, Ministry of Agriculture). Alternatively, experiments done at the SynAging site were approved by ethics committee “Comité d’Ethique Lorrain en Matière d’Expérimentation Animale”, and by the governmental agency the “Direction Départementale de la Protection des Populations de Meurthe et Moselle-Domaine Expérimentation Animale”.

### Consent for publication

All authors have approved of the contents of this manuscript and provided consent for publication.

### Availability of materials

Alpha-synuclein PFFs, the 11A5 monoclonal anti α-synuclein antibody, and the α-syn oligomers can be obtained, under MTAs, from Biogen, Prothena Biosciences, and ETAP-lab, respectively.

### Authors contributions

P.G., W.W., D.C., and M.B, designed the study. P.G., W.W., O.H., A.Ma., S.B., E.K., T.H., A.W., C.S., A. Mi., T.P., A.A., N.F. did the experiments (stereotactic surgery, tissue processing, stainings, imaging and RNA extraction). T.K., N.N. generated the microarray data. K.J.S., E.G. analyzed the microarray data. P.G., W.W., O.H., K.J.S., R.B., W.S.-S., K.B., M.M., M.B analyzed and interpreted the data. M.B. wrote the paper. All authors read and approved the final manuscript.

## Acknowledgements

We thank Laurent Vallar (Luxembourg Institute of Health) for help with gene expression arrays, Christian Jaeger (Luxembourg Centre for Systems Biomedicine) for dopamine measurements, Eliezer Masliah (University of California, San Diego) for advice, Thierry Pillot and Violette Koziel (SynAging, France) for synuclein oligomers and for advice, and Yuting Liu (Biogen) for purifying recombinant murine α-syn. We also thank Prothena Biosciences (South San Francisco, CA) for providing the 11A5 antibody.

## Supplemental figures and tables legends

**Suppl. fig.1:** Western blot characterization of α-syn moities used for intrastriatal injections. Loading of the gel was as follows: lanes 1, 6, 14 – ladder; lanes 5, 10,11- blank; lanes 2,3,4 – α-syn oligomer, 10, 100, and 500 ng respectively; lanes 7, 8, 9 – α-syn PFFs, not sonicated, 10,100, 500 ng respectively; lanes 11, 12, 13 – α-syn PFFs (sonicated), 10,100, 500 ng respectively. Bands from corresponding to the MW of α-syn monomers, dimers, and trimers are circled (red, blue, orange, respectively). Note the presence of high molecular weight moities (visible as a smear) after loading of non-sonicated PFFs. The sonication process of PFFs appears to reduce all higher molecular weight species of α-syn and the amount of dimers and monomers, since smears and monomer/dimer bands were visible after loading 100 ng of non-sonicated PFFs, but not after loading the same amount of sonicated PFFs.

**Suppl. fig. 2:** Minimal presence of α-syn inclusions in different brain regions 13 DPI after intrastriatal injection of α-syn PFFs. Only few α-syn inclusiosn were seen, ipsi- and contralaterally, in the frontal cortex, amygdala, amd ipsilaterally, but not contralaterally, in the striatum and SN. Scale bar = 130 μm.

**Suppl. Fig. 3:** No nigro-striatal degeneration and microgliosis 13 dpi after intrastriatal injection of α-syn PFFs. No loss of striatal TH-postive axons (first row), striatal DAT-positive synaptic terminals (second row), nigral TH-positive neurons (third row) was observed 13 dpi aftre injection of PFFs. No increase of Iba1-positive microglial reaction in the SN was observed (last row). Microphotographs show examples of PBS-injected control brains (ipsilateral) and ipsilateral α-syn PFF-injected brains. Scale bars = 25 μm (striatal panels), 200 μm (Subst. Nigra panels).

**Suppl. fig. 4:** Heatmaps illustrating the ventral midbrain gene expression patterns at 13 dpi and 90 dpi after intrastriatal injection of α-syn PFFs. Heatmaps are shown for p-values < 0.05 or pfp < 0.1. Scaled row expression values (row Z-score) in red indicate higher, in blue lower, and in white unchanged gene expression (see Materials and Methods for details). Comparisons of ipsi PFF *versus* ipsi PBS (upper panel row) and ipsi PBS *versus* contra PFF (lower panel row) reveal distinctive gene expression patterns at 13 dpi for both comparisons (whether higher, pfp < 0.1, or lower, p<0.05, statistical stringency is used), whereas a distinctive patterns at 90 dpi only appears in the ipsi PFF *versus* contra PFF, but not in the ipsi PFF *versus* ipsi PBS, comparisons.

## Supplemental tables legends

**Suppl table 1: Antibodies used in this study**

**Suppl table 2: Softwares used in this study**

**Suppl table 3: Molecular astrocyte and peripheral immune cell markers**

