## Supplementary figures and images for "Neurodegeneration and neuroinflammation are linked, but independent of **α**-synuclein inclusions, in a seeding/spreading mouse model of Parkinson’s disease"

### Supplemental figure 1

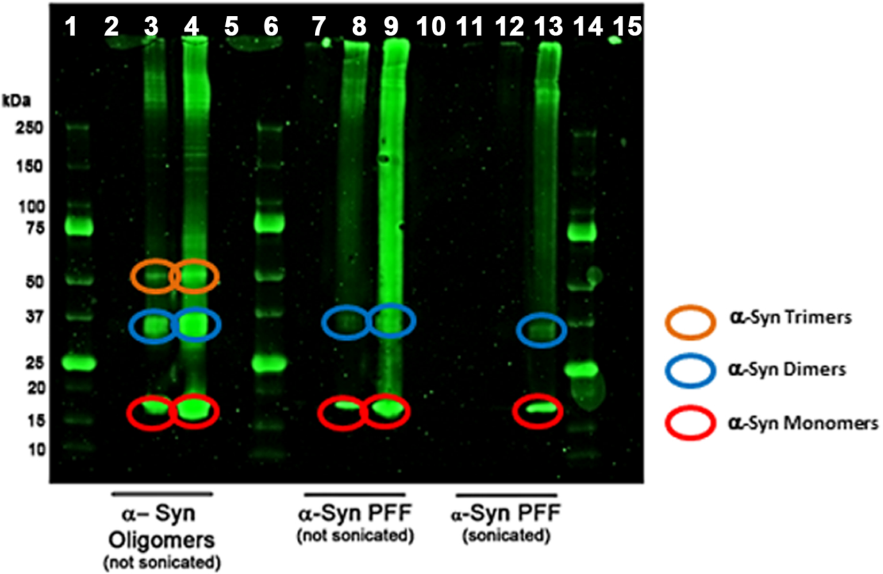

### Supplemental figure 2

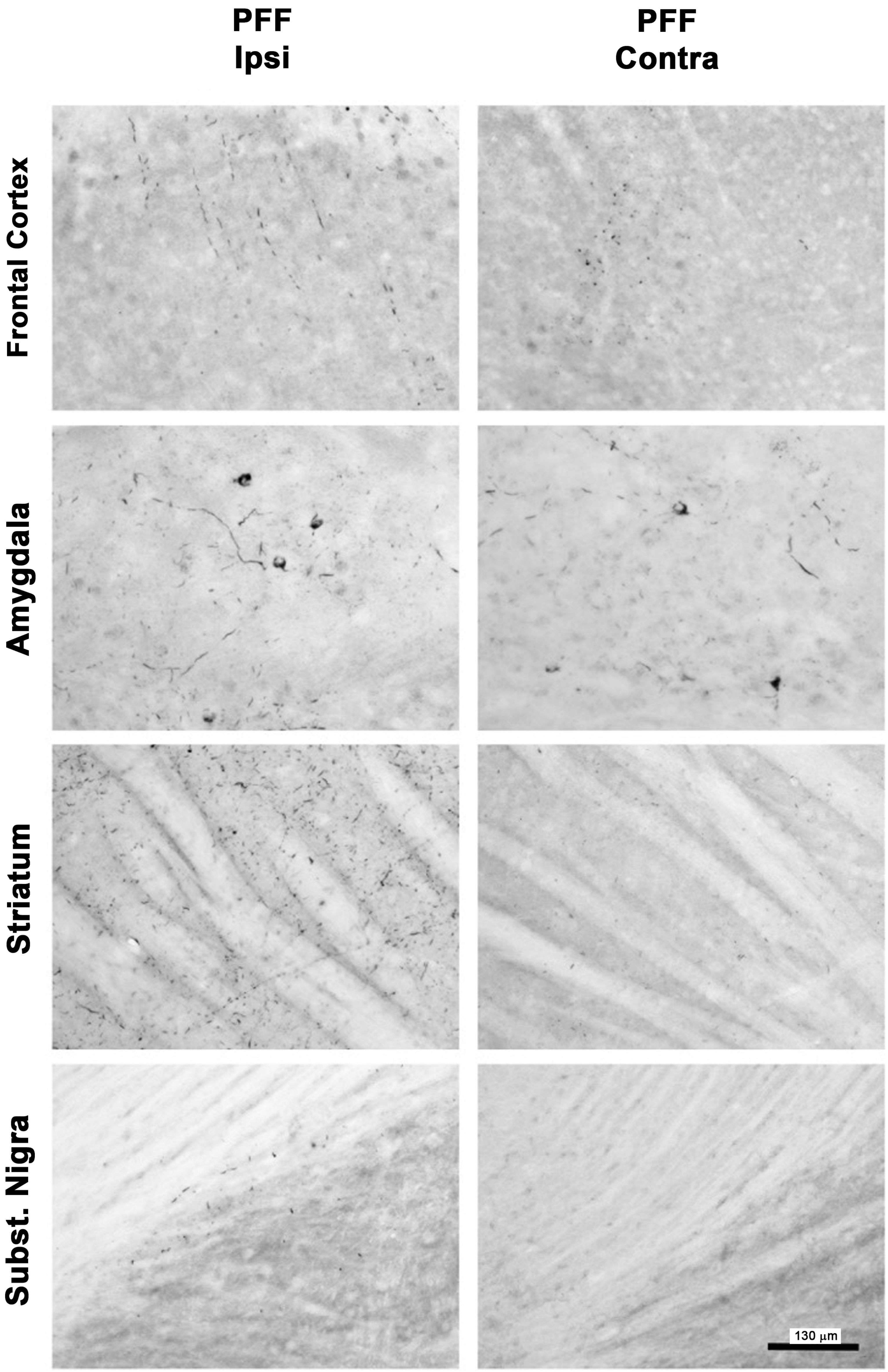

### Supplemental figure 3

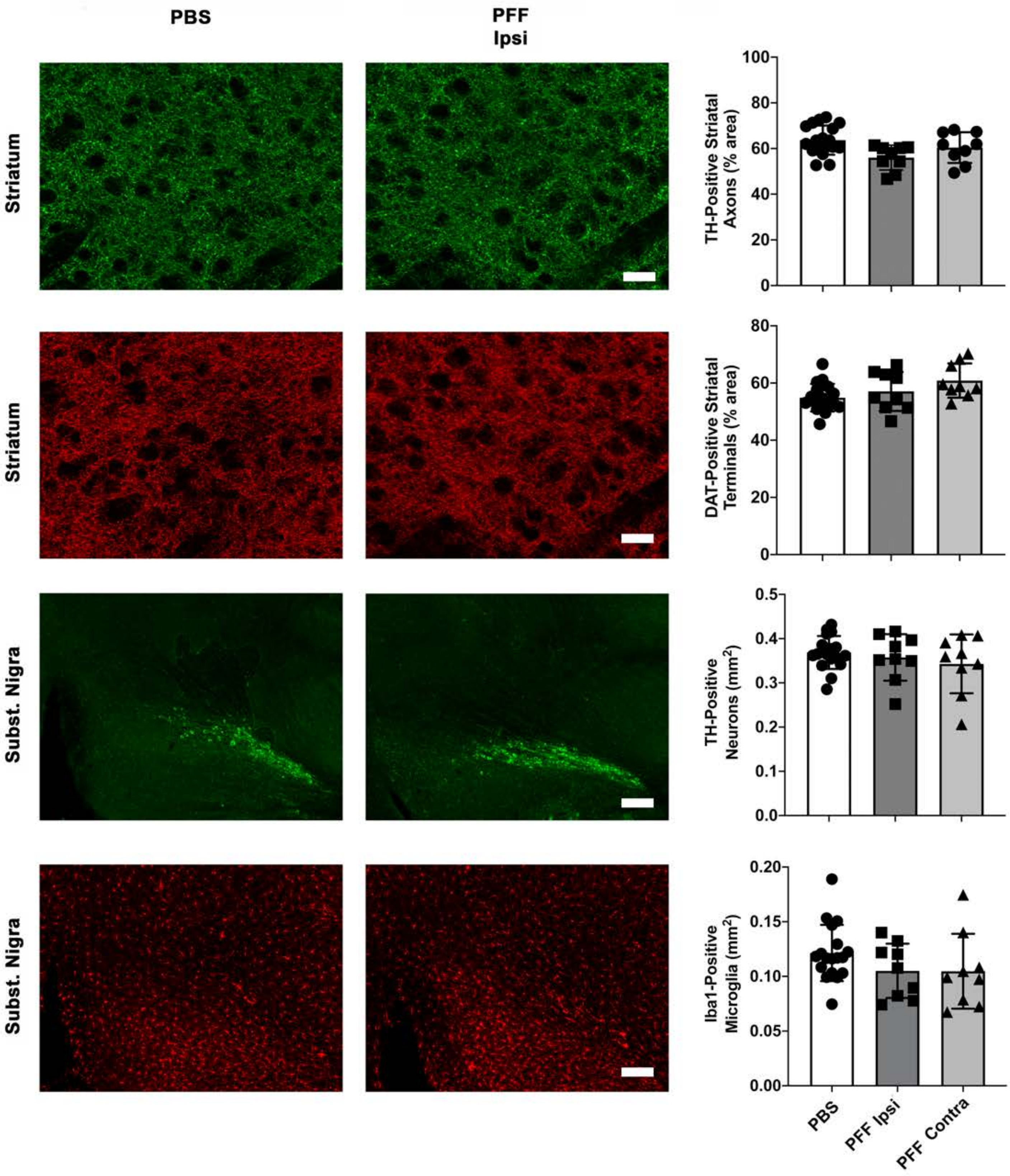

### Supplemental figure 4

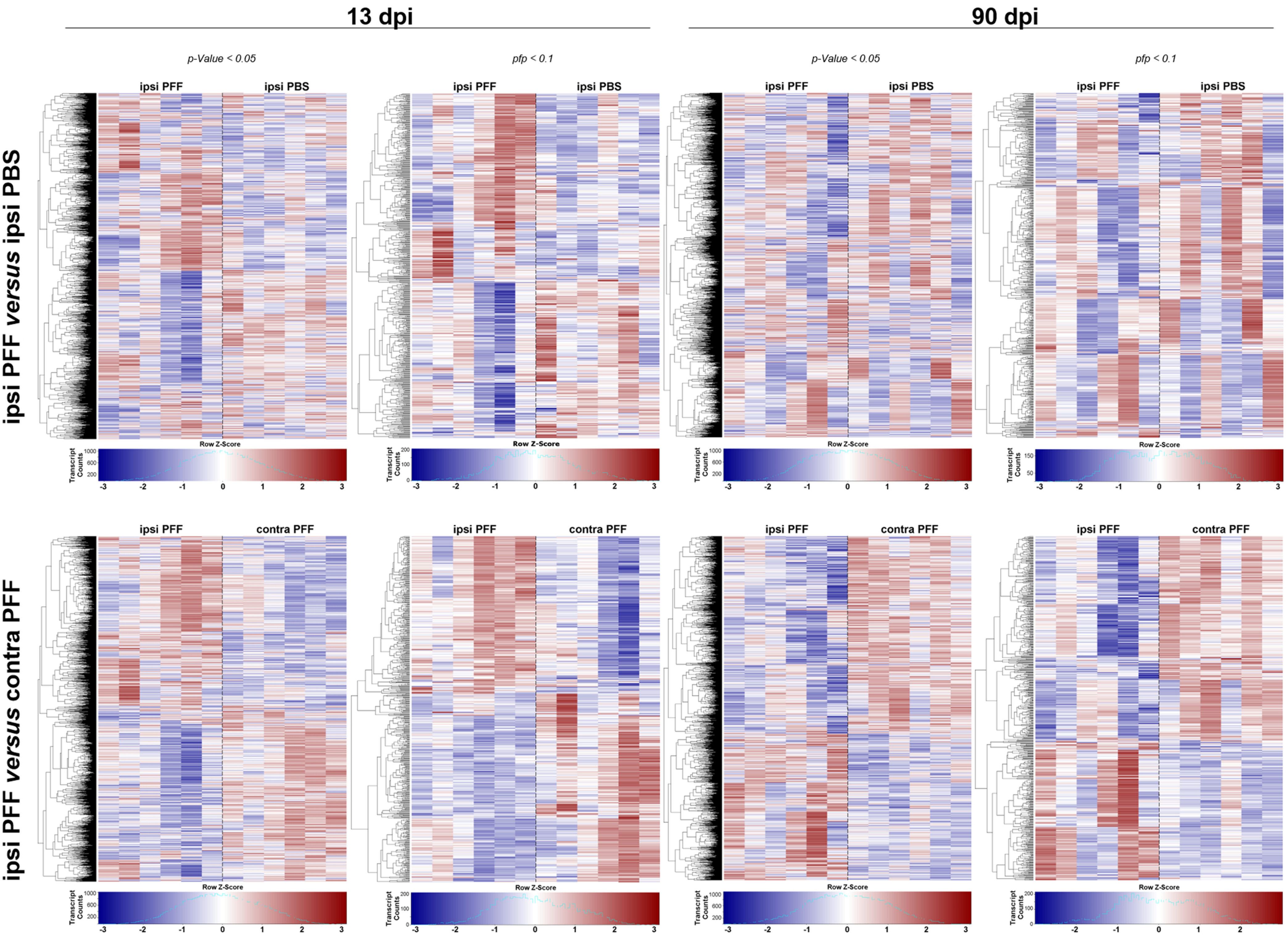
