## Supplemental table 1 for "Neurodegeneration and neuroinflammation are linked, but independent of **α**-synuclein inclusions, in a seeding/spreading mouse model of Parkinson’s disease"

| antigen | species | Manu-facturer | dilution | blocking | Secondary antibody |
|---|---|---|---|---|---|
| alpha synuclein phospho S129 | Rabbit polyclonal | Abcam Ab51253 | 1:2000 | 5% horse serum (immo-peroxidase)<br><br>5% BSA (immuno-fluorescence) | biotynilated goat-anti-rabbit (Vector Laboratories)<br><br>Alexa 488 anti-rabbit<br><br>Alexa 568 anti-rabbit |
| 11A5 Anti SER p129 α-syn | Mouse monoclonal | Prothena | n/a | n/a | PLA assay |
| ionized calcium-binding adapter molecule 1 (Iba1) | Chicken polyclonal | Abcam Ab139590 | 1:800 | 5% BSA | Alexa 568 anti-chicken |
| Iba1 | Rabbit polyclonal | Wako 019-19741 | 1:2000 | 5% BSA | Alexa 568 anti-rabbit |
| Tyrosine hydroxylase (TH) | Chicken polyclonal | Abcam Ab76442 | 1:1000 | 5% BSA | Alexa 488 anti-chicken |
| Tyrosine Hydroxylase (TH) | Rabbit polyclonal | Millipore AB152 | 1:1000 | 5% BSA | Alexa 568 anti-rabbit |
| Dopamine transporter (DAT) | Rat monoclonal | Millipore MAB369 | 1:1000 | 5% BSA | Alexa 594 anti-rat |
| Synapto-physin | Mouse monoclonal | Dakopatts Clone SY38 | 1:800 | 5% BSA | Alexa 488 anti-mouse |

Suppl table 1: Antibodies used.
