## Supplemental table 2 for "Neurodegeneration and neuroinflammation are linked, but independent of **α**-synuclein inclusions, in a seeding/spreading mouse model of Parkinson’s disease"

Suppl table 2: Software used

| Name of Software or Package | Vendor or public url | Uses |
|---|---|---|
| *PRISM 7 (v. 7.04)* | GraphPad Software Inc. (https://www.graphpad.com/) | Basic statistics (normality test, ANOVA, Kruskall-Wallis, posthoc test for multiple groups) |
| *R (v. 3.4.3)* | The R Foundation (https://www.r-project.org/) | Basic Statistical Computing and Data Manipulation |
| *R Studio (v. 1.1.414)* | https://www.rstudio.com/ | User Interface for R |
| *RankProd (v. 3.4.0)* | Bioconductor (DOI: 10.18129/B9.bioc.RankProd) | Differential Gene Expression Analysis for Microarray Datasets. A non-parametric approach. FDR statistics. |
| *VennDiagram (v. 1.6.20)* | CRAN – R (https://CRAN.R-project.org/package=VennDiagram) | Generate High-Resolution Venn and Euler Plots |
| *GSEA (v. 3.0)* | Broad Institue, Inc. (http://www.gsea-msigdb.org/gsea/index.jsp) | Analyze, annotate and interpret enrichment results |
| *Cytoscape (v. 3.6.1)* | The Cytoscape Consortium (http://manual.cytoscape.org/en/3.6.0/) | Visualizing complex networks from GSEA |
