## Supplemental table 3 for "Neurodegeneration and neuroinflammation are linked, but independent of **α**-synuclein inclusions, in a seeding/spreading mouse model of Parkinson’s disease"

| | | ipsi PFF versus ipsi PBS | | | | | | ipsi PFF versus contra PFF | | | | | |
|---|---|---|---|---|---|---|---|---|---|---|---|---|---|
| | | 13dpi | | | 90dpi | | | 13dpi | | | 90dpi | | |
| **Gene Symbols** | **Protein** | **FC** | **p-Value** | **Pfp** | **FC** | **p-Value** | **Pfp** | **FC** | **p-Value** | **Pfp** | **FC** | **p-Value** | **Pfp** |
| **ASTROCYTES** | | | | | | | | | | | | | |
| ***Gfap*** | **Glial fibrillary acidic protein** | 1.633 | **7.12E-08** | **1.85E-04** | -1.123 | **2.38E-02** | 6.34E-01 | 1.736 | **1.71E-08** | **3.62E-05** | 1.285 | **1.52E-03** | 1.29E-01 |
| ***Amigo2*** | **Adhesion molecule with Ig like domain 2** | -1.273 | 9.24E-04 | 1.73E-01 | -1.030 | 9.47E-02 | 9.12E-01 | -1.527 | **6.77E-06** | **2.86E-03** | -1.400 | **8.61E-05** | **1.51E-02** |
| ***Osmr*** | **Oncostatin M receptor** | 1.153 | **1.78E-02** | 5.87E-01 | -1.081 | **3.66E-02** | 7.10E-01 | 1.396 | **2.21E-04** | **3.30E-02** | 1.315 | **1.31E-03** | 1.16E-01 |
| ***S1pr3*** | **Sphingosine-1-phosphate receptor 3** | -1.184 | **1.71E-02** | 6.07E-01 | -1.054 | 3.00E-01 | 1.06E+00 | -1.159 | 7.02E-02 | 8.12E-01 | -1.118 | 1.19E-01 | 8.61E-01 |
| ***Vim*** | **Vimentin** | 1.183 | **2.82E-02** | 7.02E-01 | -1.246 | **6.29E-03** | 4.18E-01 | 1.135 | 9.37E-02 | 8.74E-01 | 1.147 | 6.38E-02 | 7.42E-01 |
| ***Serping1*** | **Serine/ cysteine peptidase inhibitor, member 1** | -1.031 | 1.05E-01 | 9.94E-01 | -1.513 | **1.35E-05** | **2.85E-02** | 1.223 | **6.77E-03** | 2.65E-01 | 1.764 | **1.40E-07** | **1.82E-04** |
| ***Ptx3*** | Pentraxin related gene | -1.028 | 5.39E-01 | 1.04E+00 | 1.023 | 6.50E-01 | 9.97E-01 | -1.000 | 6.48E-01 | 1.06E+00 | 1.025 | 5.57E-01 | 1.06E+00 |
| **PERIPHERAL IMMUNE CELLS** | | | | | | | | | | | | | |
| ***Ptprc*** | **Protein tyrosine phosphatase, receptor C (CD45)** | 1.457 | **1.69E-05** | 1.02E-02 | -1.063 | 2.00E-01 | 1.03E+00 | 1.516 | **2.72E-05** | **7.71E-03** | 1.193 | **3.05E-02** | 5.87E-01 |
| ***Cd19*** | CD19 antigen | 1.004 | 6.65E-01 | 1.02E+00 | 1.039 | 4.75E-01 | 1.02E+00 | -1.054 | 4.60E-01 | 1.08E+00 | 1.024 | 5.14E-01 | 1.06E+00 |
| ***Cd3e*** | CD3 antigen, epsilon polypeptide | 1.037 | 4.32E-01 | 1.07E+00 | 1.045 | 2.82E-01 | 1.06E+00 | -1.009 | 5.92E-01 | 1.07E+00 | 1.037 | 4.05E-01 | 1.03E+00 |
| ***Cd4*** | **CD4 antigen** | 1.172 | **3.63E-02** | 7.59E-01 | -1.105 | **1.63E-02** | 5.52E-01 | 1.094 | 7.61E-02 | 8.15E-01 | 1.243 | **4.32E-03** | 2.30E-01 |
| ***Cd8a*** | CD8 antigen | -1.003 | 4.77E-01 | 1.06E+00 | -1.141 | 9.30E-02 | 9.07E-01 | 1.046 | 2.44E-01 | 1.06E+00 | 1.076 | 3.19E-01 | 1.01E+00 |
